# Improving racial fairness in brain age models using style-transfer synthesis

**DOI:** 10.64898/2026.09.16.751796

**Authors:** Vittoria Banchieri, Dina Zemlyanker, Juan Eugenio Iglesias, the Alzheimer’s Disease Neuroimaging Initiative

## Abstract

Brain age models can systematically mispredict age for specific demographic groups, risking biased estimates of neurological health. We present an approach to improve accuracy and reduce racial disparities using synthetic T1 images generated via style-transfer with SuperSynth, an open-source FreeSurfer tool that produces intensity harmonized isotropic images regardless of the input’s contrast or resolution. We refer to the original scans as the real domain and their SuperSynth-derived counterparts as the synthetic domain. Because each synthetic image derives from the participant’s own scan, comparisons between domains hold anatomy and demographic composition constant. We audited fairness by training a neural network across thirteen racial compositions on a diverse cohort (683 White, 605 Black, 431 Asian) in both domains. Our findings highlight three key insights. First, the synthetic domain enhanced both fairness and accuracy, and even models trained on a single demographic showed reduced racial disparity purely from switching domain. The same pattern appeared in two independently developed pretrained models. Second, models trained on synthetic data demonstrated superior out-of-distribution robustness in external clinical testing. Third, augmenting imbalanced datasets with synthetic minority images closed 61% of the fairness gap without requiring new data collection, and unlike explicitly supplying race labels, does not require race as a model input at deployment. Representational analyses indicated that appearance standardization changes which features drive age prediction rather than removing demographic information from the images. By utilizing synthetic data, our approach offers a robust pathway to more equitable and generalizable brain age models.

## 1 INTRODUCTION

Since the seminal work from Franke et al. (2010) showed that age can be predicted from structural brain magnetic resonance imaging (MRI) using computational methods, the discrepancy between predicted and real age, known as BAG (brain age gap), has become a widely used index of brain health, associated with cognitive decline, neurodegeneration, and mortality risk (Cole and Franke 2017, Cole et al. 2018). For example, each one-year increase in BAG corresponds to a increased risk of Alzheimer’s disease (AD) by 16.5%, mild cognitive impairment (MCI) by 4.0%, and all-cause mortality by 12% (Zhang et al. 2025); if a model systematically mispredicts for a particular demographic group, it does not merely make larger errors, but misclassifies the neurological risk of that group. For brain age to fulfill its clinical promise, robustness and fairness across demographic groups must be ensured (Ricci Lara et al. 2022).

The best-performing brain age models are currently neural network (NN)-based, reaching mean absolute errors (MAE) below 3 years (La Rosa et al. 2025), reflecting the broader success of deep learning in medical imaging (Litjens et al. 2017, Shen et al. 2017). As these models become increasingly accurate and clinically relevant, concerns about their fairness have intensified. As established in the literature (Ricci Lara et al. 2022, Xu et al. 2024), we consider a model unfair if it systematically advantages one demographic subgroup over another; in this study, the subgroups of interest are self-reported race and biological sex. Consistent with this concern, demographic performance disparities have been documented across a range of medical imaging tasks, including cardiac MRI segmentation (Puyol-Antón et al. 2022), chest radiograph classification (Seyyed-Kalantari et al. 2021), brain MRI segmentation (Ioannou et al. 2022, Danaee et al. 2025), brain MRI reconstruction (Du et al. 2023), and prostate MRI segmentation (Alqarni et al. 2025).

A possible source for disparities in model performance is shortcut learning. Convolutional neural-networks (CNNs) have been shown to predict protected attributes (characteristics that can be used to discriminate individuals (Zhang and Wu 2017), such as race, sex, and age) directly from medical images (Gichoya et al. 2022, Duffy et al. 2022, Adleberg et al. 2022, Yang et al. 2024). This capability raises concern that models may exploit demographic or acquisition-related image features rather than the underlying biological signal of interest (DeGrave et al. 2021, Banerjee et al. 2023, Geirhos et al. 2020, Brown et al. 2023). This shortcut learning can produce systematic performance disparities even without access to demographic labels (Geirhos et al. 2020) and is increasingly recognized as a major obstacle to trustworthy and equitable medical imaging artificial intelligence (Geirhos et al. 2020, Xu et al. 2024, Wang et al. 2024).

Demographic disparities in brain age models are well documented, yet the factors underlying them remain difficult to disentangle. Training composition and sample size are consistently confounded: most studies evaluate models trained on naturally White-skewed cohorts without varying racial composition at fixed sample size (Piçarra and Glocker 2023, Dempsey et al. 2023, Adkins and Hanson 2025), and even studies that rebalance cohorts through upsampling typically do not independently control for sample size (Gravina et al. 2026). Age bias correction introduces an additional source of variability, as it is applied inconsistently or omitted entirely, despite the well-established tendency of brain age models to exhibit regression-to-the-mean effects that can distort group comparisons (de Lange et al. 2022, Beheshti et al. 2019, Piçarra and Glocker 2023, Dempsey et al. 2023, Adkins and Hanson 2025, Aguzin Parrilli and Belzunce 2026); some studies adjust for age statistically rather than correcting predictions directly (Gravina et al. 2026). Comparisons across multiple pretrained models further complicate interpretation because differences in architecture, training data, preprocessing, and bias-correction procedures cannot be separated (Aguzin Parrilli and Belzunce 2026, Adkins and Hanson 2025). Consequently, it remains unclear to what extent reported disparities arise from model design, data imbalance, methodological choices, or underlying biological variation.

A potential avenue for mitigating these performance gaps is the use of synthetic MRI. In medical imaging, synthetic data have been explored both as a data augmentation strategy and as a means of reducing acquisition-related variability. As augmentation, synthetic images can supplement sparsely sampled groups and improve robustness (Jackson et al. 2025, van Breugel et al. 2024 2023ba 2021). More recently, synthesis methods have also been used to mitigate differences arising from scanner hardware, imaging sites, and acquisition protocols (Dewey et al. 2019, Zuo et al. 2021). Among these, style transfer approaches define appearance standardization as an image-to-image problem, re-rendering each scan in a standardized appearance rather than generating new subjects (Liu et al. 2023).

A particularly promising way of training such models is domain randomization, a technique in which neural networks are exposed to images with continuously randomized contrast, resolution, orientation, and noise characteristics at every training iteration (Billot et al. 2023ab, Hoffmann 2025, Iglesias et al. 2023). By experiencing an effectively unlimited range of acquisition conditions during training, the network is discouraged from relying on protocol-specific image features (image contrast or resolution) and instead learns anatomy-driven representations that generalize across imaging environments. This principle underlies SynthSR and its successor SuperSynth (Billot et al. 2023a, Iglesias et al. 2023), both available through FreeSurfer (Fischl 2012). These style transfer models transform heterogeneous clinical MRI scans into 1 mm isotropic T1s, harmonized in terms of intensities and resolution, without requiring paired data or scanner labels. Throughout this work we refer to the original acquisitions as the real domain and their SuperSynth-derived counterparts as the synthetic domain. Domain randomization has also been applied directly to brain age prediction through SynthBA (Puglisi et al. 2024, Biondo et al. 2025), which was designed to be robust to acquisition variability. Importantly, using style transfer derived synthetic scans does not appear to come at the expense of accuracy: a CNN retrained on super-resolved SynthSR-derived images achieved brain age prediction performance comparable to the same model trained on native scans across multiple clinical MRI modalities (Valdes-Hernandez et al. 2023).

This work makes four contributions:

- We show that style-transfer synthesis improves accuracy and reduces racial bias. This is established by training and evaluating the same brain age model on both real T1-weighted scans and their SuperSynth-derived synthetic counterparts, holding participant anatomy, validation and test sets, and evaluation procedures constant. We assess the robustness of this effect across thirteen racial training compositions at matched sample sizes, using a fixed architecture (Peng et al. 2021) and a uniform age bias correction procedure, which separates the contribution of imaging domain from that of training composition and sample size. We further test whether the effect generalizes beyond our own architecture by evaluating BrainAgeNeXt (La Rosa et al. 2025) and SynthBA (Puglisi et al. 2024, Biondo et al. 2025) on the same fixed test set.
- We demonstrate that style-transfer synthesis improves generalization. We evaluate out-of-distribution performance on an ADNI3 (Alzheimer’s Disease Neuroimaging Initiative) cohort (Jack et al. 2008, Weiner et al. 2017), a population differing from the training data in acquisition environment, age distribution, and clinical composition, and assess whether improved transfer preserves the clinical sensitivity of the brain age gap across diagnostic groups.
- We show that synthetic data can also be used selectively, as a targeted mitigation strategy. We compare synthetic oversampling of minority subjects against race-label injection (Gravina et al. 2026) in a White-dominant training set, and find that combining real and synthetic images outperforms either the fully real or the fully synthetic version of the same configuration. Unlike race-label injection, oversampling uses demographic information only during dataset construction and does not require race as a model input at deployment.
- We characterize the mechanisms underlying performance and fairness differences using t-distributed stochastic neighbor embedding (t-SNE) (van der Maaten and Hinton 2008), linear probing (Alain and Bengio 2016), and integrated gradients (IG) (Sundararajan et al. 2017), characterizing how age and demographic information are represented within the models.

## 2 METHODS

### 2.1 MRI data

#### MGH Dataset

We used a subset of the cohort described by Liu et al. (2025), drawn from the Massachusetts General Hospital (MGH) clinical imaging archive via the Research Patient Data Registry: the full cohort comprises N=5,977 subjects aged 18–90, free of documented structural brain abnormality such as tumor, stroke, or traumatic brain injury, and with complete self-reported race (Asian, Black, or White). Because these are clinical rather than research acquisitions, sequence type is not reliably recorded, so T1-weighted scans were identified using heuristics on the images as follows. For each scan we ran SynthSeg (Billot et al. 2023a) to extract median white-matter (WM), gray-matter (GM), and cerebrospinal fluid (CSF) intensities and identified T1-weighted-like scans as those exhibiting the expected intensity ordering (WM > GM > CSF). We retained every subject with at least one such scan at near-1 mm isotropic resolution, the input expected by most neuroimaging tools and published brain age models, so that the cohort could be processed directly and results compared across methods. Applying these criteria yielded a final cohort of N=1,719 (803 male; 683 White, 605 Black, 431 Asian). Finally, for subjects who had multiple T1 scans available, a single one was randomly selected to avoid the same individual contributing multiple times to training or evaluation cohorts.

#### ADNI Dataset

To probe the generalization ability of the best-performing models identified in this study, we further evaluated them on 1 mm isotropic T1-weighted scans from 302 ADNI3 participants (Jack et al. 2008), comprising patients with mild cognitive impairment (MCI, n=132), subjects reporting subjective memory complaints (SMC, n=49) and cognitively normal controls (CN, n=121). As in the MGH cohort, we considered only participants aged 90 or below; the resulting ADNI3 sample ranged from 61 to 90 years.

The ADNI was launched in 2003 as a public-private partnership, led by Principal Investigator Michael W. Weiner, MD. The primary goal of ADNI has been to test whether serial magnetic resonance imaging (MRI), positron emission tomography (PET), other biological markers, and clinical and neuropsychological assessment can be combined to measure the progression of MCI and early AD.

### 2.2 Image preprocessing

One T1-weighted scan per subject (selected as described above for the MGH cohort, and the single available scan for ADNI3) was processed with SuperSynth (Iglesias et al. 2023, Billot et al. 2023a), which simultaneously (i) resampled the input to 1*×*1*×*1 *mm*^3^ isotropic resolution, (ii) generated a 1*×*1*×*1 *mm*^3^ contrast-normalized synthetic T1-weighted image, (iii) produced an affine transformation file, and (iv) derived segmentation labels. Both the resampled real and the synthetic T1 were intensity-normalized to [0, 1] and resampled to the space of the MNI152 1 mm isotropic template using the SuperSynth-derived affine transform. As introduced above, we refer to the registered resampled T1-weighted scan as the *real domain* and the SuperSynth-derived image as the *synthetic domain*. Because both images derive from the same subject anatomy, this pipeline enables within-subject paired comparisons across imaging domains while holding all anatomical and demographic variables constant.

### 2.3 Model architecture and training

The architecture employed is a Simple Fully Convolutional Network (SFCN) (Peng et al. 2021), selected because it combines state-of-the-art brain age prediction performance with strong test–retest reliability (Peng et al. 2021, Dörfel et al. 2023); a schematic representation of the architecture is shown in Figure 1. GroupNorm (8 groups per layer) (Wu and He 2018) was used in place of BatchNorm3d (Ioffe and Szegedy 2015) throughout, as BatchNorm produced unstable training in smaller-sample subsets of the composition sweep (Supplementary Materials Section 1). All models were trained with Adam (lr = 3*×*10^−5^, batch size = 4) to minimize the mean squared error between predicted and chronological age, with early stopping on validation MAE (patience = 10) and a maximum of 100 epochs. Following the resampling to the MNI template space, the input was a 3D volume of 193 *×* 229 *×* 193 voxels at 1 mm isotropic resolution. To account for variability due to random initialization and stochastic optimization, each model configuration was trained independently five times on the same data and the results aggregated by averaging.

**FIGURE 1.**
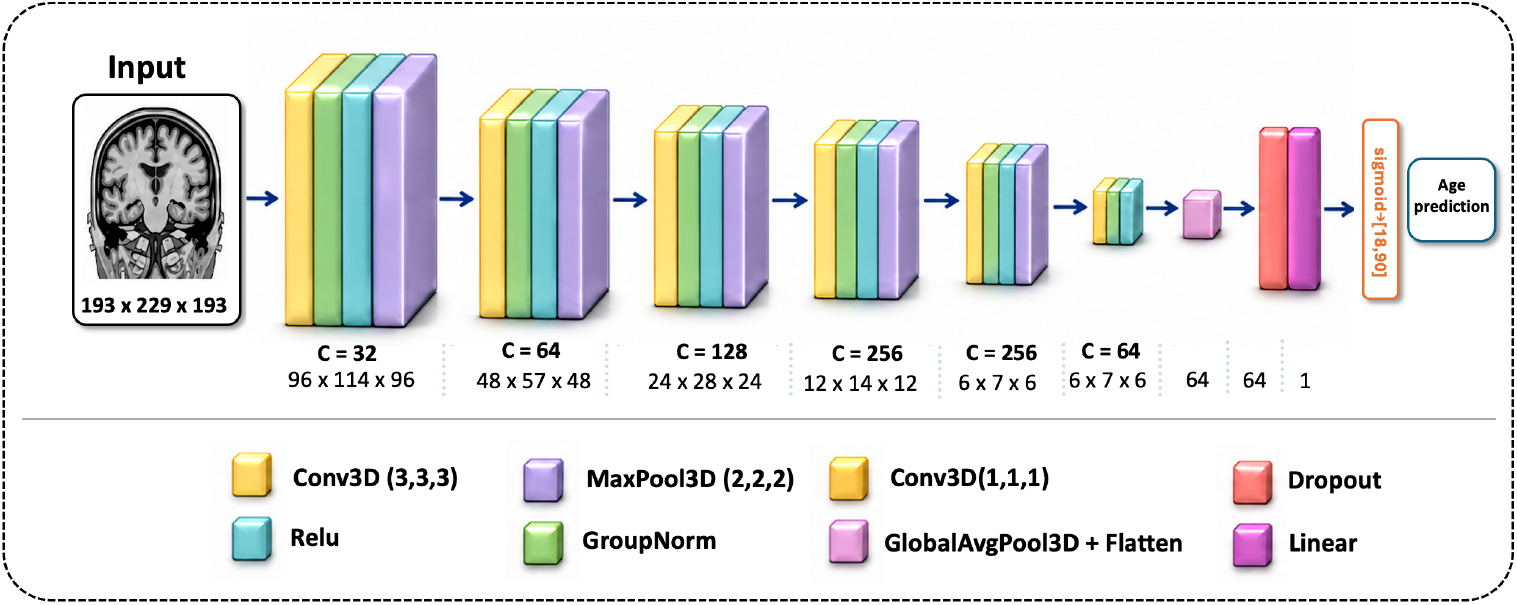
Brain age prediction architecture. The SFCN-based convolutional neural network used throughout this study processes three-dimensional MRI volumes through successive convolutional blocks and spatial downsampling layers to learn hierarchical anatomical features. These features are then globally aggregated and passed to a final prediction layer to estimate brain age. The architecture and training procedure were held fixed across experiments; only the training set composition and imaging domain were varied. C denotes the number of channels.

### 2.4 Age bias correction

Let ŷ_*i*_ be the brain age of a subject *i* in a dataset, predicted from their MRI scan, and let *y*_*i*_ be their chronological age at the time of the scan. The BAG is then defined as:

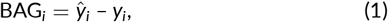

Brain age models systematically over-predict age for younger subjects and under-predict for older subjects, a well-documented phenomenon known as age bias (de Lange et al. 2022, Beheshti et al. 2019, Smith et al. 2019, Franke et al. 2010, Cole and Franke 2017, Zhang et al. 2023, Liang et al. 2019, Treder et al. 2021), arising from the statistical tendency of any regression model trained on a bounded age range to predict values closer to the training mean. This manifests as a significant negative correlation between BAG and chronological age. Following standard practice (Zhang et al. 2023), we applied a uniform two-stage correction to our aggregated predictions.

- **Step 1: Linear (Beheshti) correction**. A linear model was fitted relating BAG to chronological age on the averaged validation set predictions:

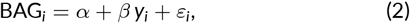

where *α* and *β* are estimated by ordinary least squares on the validation set. The fitted coefficients (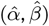) were then applied to the averaged test set predictions to obtain the Beheshti-corrected BAG (Beheshti et al. 2019):

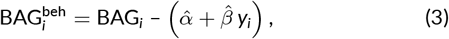

and the corresponding Beheshti-corrected predicted age as 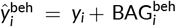. This removes the bulk of the sample-level bias.
- **Step 2: Soft-Gaussian age-bin correction**. A residual, age-structured bias often remains after Step 1, concentrated at the extremes of the age range. This was corrected in two parts, both computed on the test set. First, test subjects were partitioned into four fixed age bins (18–37, 38–57, 58–77, 78–90 years), and the mean Beheshti-corrected BAG was computed within each bin:

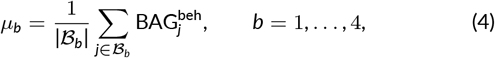

where *B*_*b*_ is the set of test subjects whose age falls in bin *b*, and each bin mean *µ*_*b*_ is associated with that bin’s midpoint *c*_*b*_. Second, each individual subject’s correction was computed as a Gaussian-weighted interpolation across these four bin means, rather than a hard assignment to their own bin:

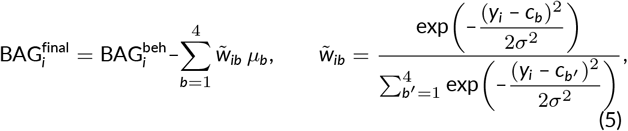

with *σ* = 5 years. This softens the hard bin boundaries: a subject near a bin edge draws on a blend of the neighboring bins’ mean residuals rather than only their own bin’s mean, avoiding discontinuities at bin edges while still removing the bulk of the residual age-structured bias. This drives any residual age-structured BAG bias towards zero. The necessity of a second correction step beyond simple linear debiasing has been demonstrated empirically by Zhang et al. (2023). Step 1 is therefore the deployable correction (parameters estimated on validation data, applied to held-out test data), while Step 2 requires test set statistics. The full pipeline of correction is shown in Supplementary Materials, Section 2.

### 2.5 Performance and fairness metrics

Each model’s performance was assessed using the mean absolute error between predicted age (ŷ_*i*_) and chronological age (*y*_*i*_):

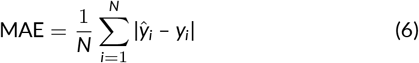

and the coefficient of determination:

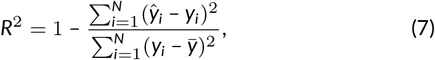

where 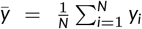 is the mean chronological age. *R*^2^ was computed on uncorrected predictions as the primary measure of intrinsic model accuracy; MAE was reported after each correction stage, with the Beheshti-corrected value as the primary accuracy metric. The racial fairness gap (RFG) computed also at each correction stage was our primary fairness outcome:

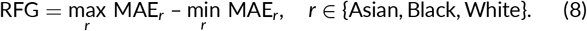

RFG was additionally computed within subgroups defined by sex and race to complete the demographic audit.

To verify that each correction stage achieved its intended goal, Pearson correlations between BAG and chronological age were computed and tested for significance at each stage, for each of the twenty-six combinations given by the thirteen training configurations evaluated in the two domains. Within-model effects on BAG were assessed using Kruskal–Wallis tests (Kruskal and Wallis 1952) for race, for sex, and for subgroups defined jointly by sex and race, with effect sizes reported as epsilon-squared. Because each test set contained only 44 subjects per racial group, these within-model subgroup analyses were considered exploratory and used only to characterize potential demographic effects. Formal statistical comparisons were instead performed on model-level fairness gaps, treating each training configuration as one observation. Racial and sex fairness gaps, and the effect of imaging domain on the racial fairness gap, were compared across the training configurations using Wilcoxon signed-rank tests (Wilcoxon 1945), treating each configuration’s RFG as one paired observation. Throughout, p-values were false discovery rate (FDR)-corrected within each test family using the Benjamini–Hochberg procedure (Benjamini and Hochberg 1995). All analyses were conducted in Python using SciPy (Virtanen et al. 2020) and scikit-learn (Pedregosa et al. 2011).

## 3 EXPERIMENTS AND RESULTS

### 3.1 Experimental design

Using simultaneous stratification by race, sex, and twenty-year age bins, subjects from the MGH Dataset were divided into non-overlapping training, validation, and test sets. An overview of the cohort partitioning and training configurations is shown in Figure 2.

**FIGURE 2.**
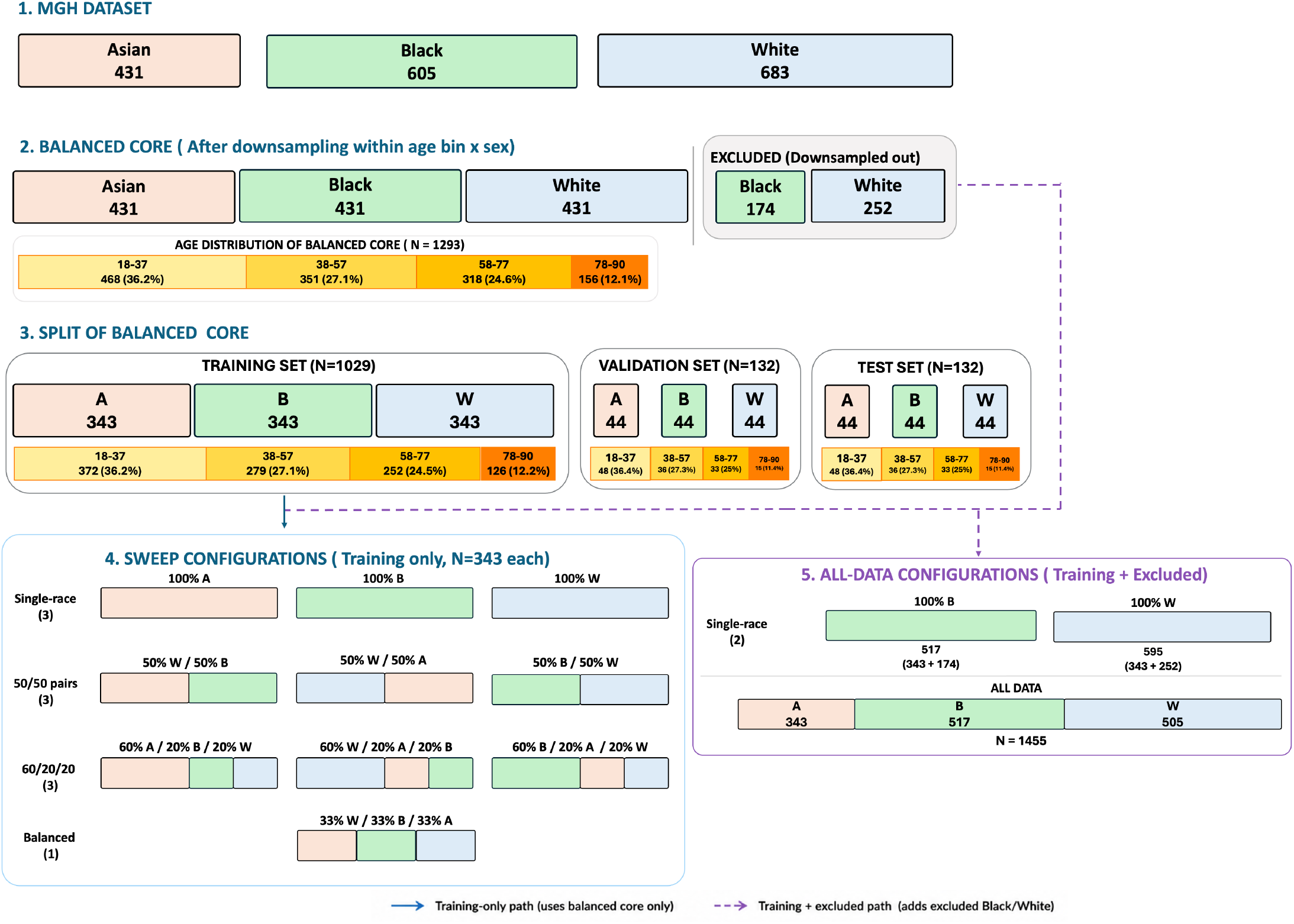
Overview of the MGH cohort and the thirteen training-composition configurations. The full racially balanced pool (*N* = 1,719; 683 White, 605 Black, 431 Asian) was split into fixed, non-overlapping training, validation, and test sets, with validation and test sets held constant at 132 subjects each (44 per race, matched by sex and age) across all experiments. From the remaining training pool, thirteen configurations were constructed: the all-data model, two full single-race models, three fixed-size (*n* ~ 343) single-race models, three pairwise 50/50 configurations, one balanced 33/33/33 configuration, and three imbalanced dominant-group configurations.

The validation and test sets each contained 132 subjects: 44 Asian, 44 Black, and 44 White matched by sex and age distribution. These sets were held fixed across all experiments, ensuring that differences between models reflected changes in the training data or imaging domain. The remaining subjects (*n* = 1,455) formed the training pool from which thirteen configurations were constructed. These included the complete training dataset (*n* = 1,455), two full single-race datasets (White = 595, Black = 517), and ten fixed-size (*n* = 343) datasets: three single-race, three pairwise 50/50, one balanced 33/33/33, and three dominant-group datasets containing either 60% White 20% Asian 20% Black, 60% Black 20% Asian 20% White or 60% Asian 20% White 20% Black subjects. Fixed-size configurations were capped at 343 subjects, because this was the size of the smallest racial group. This allowed us to vary training composition while holding total sample size constant. Full single-race datasets were additionally compared with their fixed-size counterparts to assess the separate contribution of training set size.

We assessed each configuration in the two domains described above. Models in the real domain were trained and evaluated using the real T1 acquisitions, whereas models in the synthetic domain used the corresponding SuperSynth-derived T1 images. The same subjects, architecture, optimization procedure, validation set, and test set were used for matched real–synthetic comparisons. Unless otherwise stated, reported MAE and RFG values refer to predictions corrected using the linear procedure. Complete participant counts and results for all configurations are reported in Section 3 of Supplementary Materials.

#### Age bias correction

Beheshti-corrected values are treated as primary results, since that stage is estimated on validation data and is therefore deployable without knowledge of test set statistics. Fully corrected values are reported alongside them to confirm that findings persist after complete removal of age-structured bias. Both stages were necessary: a significant residual BAG–age correlation remained after linear correction in 10 of 13 real domain and 9 of 13 synthetic domain configurations, and was reduced to non-significance in all 26 combinations only after the soft-bin step. Full per-configuration correlations are reported in Section 2 of Supplementary Materials.

### 3.1 Existing pretrained models exhibit racial bias

To assess whether state-of-the-art pretrained models show performance disparities among racial subgroups, we evaluated BrainAgeNeXt (La Rosa et al. 2025) and SynthBA (Puglisi et al. 2024, Biondo et al. 2025) on our fixed, racially balanced MGH real domain test set. Both models require skull-stripped inputs, so the preprocessing of Section 2.2 was followed by skull-stripping with the corresponding SuperSynth segmentation; predictions were then evaluated using the age bias correction procedure and fairness metrics described in Methods. Full per-race and correction-stage results are reported in Supplementary Materials, Section 4.

BrainAgeNeXt achieved an uncorrected *R*^2^ = 0.905 and a Beheshti-corrected MAE of 4.76 *±* 3.70 years, with an RFG of 1.30 years. Errors were unevenly distributed across racial groups: Black subjects had the highest MAE at 5.45 years, against 4.66 for White and 4.15 for Asian subjects; this ordering persisted after the soft-bin correction, which narrowed the RFG to 0.82 years.

SynthBA showed larger disparities. Uncorrected *R*^2^ was 0.487 for White subjects but only 0.197 and 0.129 for Black and Asian subjects, indicating a pronounced White performance advantage. Overall performance was also substantially lower, with an uncorrected MAE of 14.32 *±* 10.34 years against 5.07 *±* 3.91 for BrainAgeNeXt, and an uncorrected RFG of 4.13. Because *R*^2^ was low in the non-White subgroups, the linear correction was fitted to uninformative predictions and may therefore absorb noise rather than systematic age bias. The corrected metrics should consequently be interpreted cautiously: MAE decreased to 8.52 years and RFG to 1.33 years after Beheshti correction, and to 8.23 and 0.96 years after full correction.

### 3.3 Style-transfer synthesis improves accuracy and reduces racial bias

Matched real and synthetic domain configurations were compared to isolate the effect of synthetic standardization while holding participant anatomy, training composition, model architecture, and evaluation data constant (Table 1). Synthetic domain models showed significantly smaller racial fairness gaps at both correction stages. After Beheshti correction, mean RFG decreased from 2.12 years in the real domain to 1.43 years in the synthetic domain, with improvement in 10 of 13 matched configurations (paired Wilcoxon, n=13 model pairs, p=0.013). After the full correction protocol RFG was lower in all 13 synthetic domain configurations (mean 1.33 versus 2.19 years, p=0.0002).

**TABLE 1.**
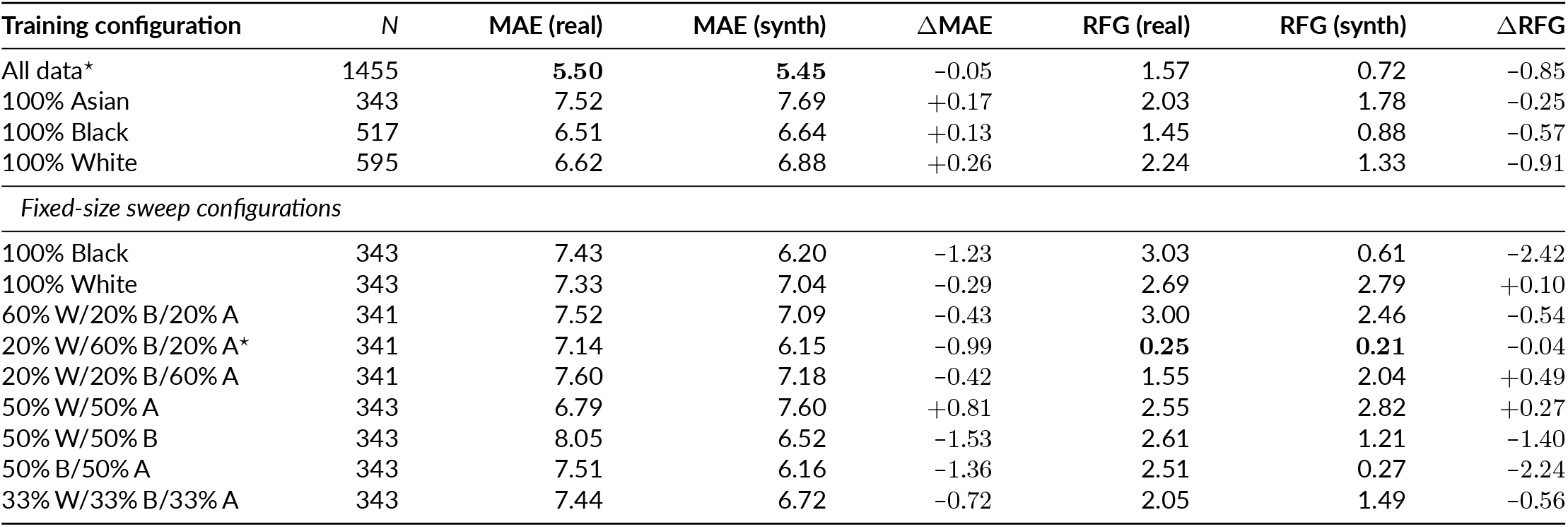
MGH Dataset performance across the thirteen training configurations. Overall MAE and RFG are reported after Beheshti correction on the fixed balanced test set. Differences are calculated as synthetic minus real, so negative values indicate improvement in the synthetic domain. Bold values indicate the best result in each metric column. The star marks configurations whose synthetic domain model was Pareto-optimal across MAE and RFG. MAE and RFG are expressed in years.

Importantly, this did not come at the cost of accuracy. Overall MAE was lower in the synthetic domain in 9 of 13 configurations after linear correction and in 12 of 13 after full soft-binning correction. Uncorrected *R*^2^ was higher in the synthetic domain for most configurations and improved on average for every racial group, increasing from 0.62 to 0.73 for Asian subjects, 0.65 to 0.70 for Black subjects, and 0.76 to 0.81 for White subjects.

In the real domain, fixed-size RFG ranged from 0.25 to 3.03 years. Training with synthetic scans reduced disparities across much of this range and did not need demographic training set diversity to be effective. Among the five single-race configurations, RFG decreased in four after switching to the synthetic domain, including the fixed-size (*n* = 343) Black-only model (3.03 to 0.61 years) and the full (*n* = 595) White-only model (2.24 to 1.33 years). The fixed-size White-only model was the sole exception. Heavily White-dominant training also retained a comparatively large synthetic domain gap, indicating that image standardization largely attenuates but does not completely eliminate the influence of training composition.

Pareto analysis of overall MAE and RFG identified only synthetic domain configurations as jointly optimal (Figure 3). The all-data synthetic model provided the lowest MAE (MAE 5.45 years; RFG 0.72 years), whereas the synthetic 20% W/60% B/20% A model provided the lowest RFG (MAE 6.15 years; RFG 0.21 years). Every real domain configuration was dominated on both objectives by at least one synthetic domain configuration.

**FIGURE 3.**
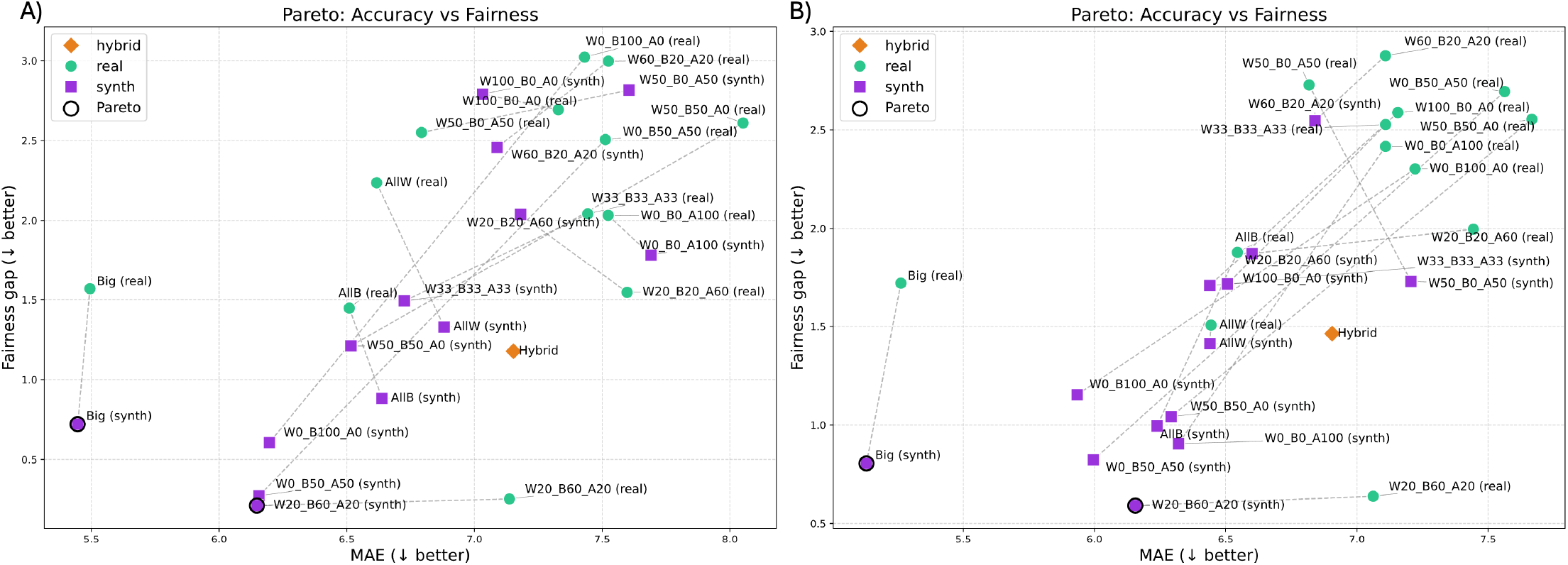
Each point is a training configuration evaluated on the fixed 132-subject test set, plotted by overall MAE (x-axis) against the racial fairness gap (y-axis); lower is better on both axes. A) Results after linear correction. B) Results after linear correction + soft binning. The color green denotes models tested in the Real MRI domain, the color purple in the Synthetic T1 domain, and the orange diamond marks the Hybrid configuration (60W/20B/20A real training augmented with SynthT1 minority scans, evaluated on real validation). Dashed lines connect matched real-synth pairs of the same training composition, visualizing the effect of domain switch on each configuration.

Race gaps exceeded sex gaps in 12 of 13 real-domain configurations (paired Wilcoxon, p=0.0017). Switching to the synthetic domain reduced the mean linear-corrected race gap from 2.12 to 1.43 years (32%), whereas the sex gap did not decrease (1.07 to 1.27 years, *p* = 0.068); race and sex disparities consequently converged and no longer differed significantly in the synthetic domain (*p* = 0.64). Within-model sex effects were small (Kruskal–Wallis, *ϵ*^2^ = 0.000 – 0.030) and did not survive FDR correction in any configuration. Asian Male exhibited the largest MAE in 7 of 13 real domain configurations, whereas Black Female subjects exhibited the largest MAE in all 13 synthetic domain configurations (MAE 6.26–9.88 years), a subgroup whose disadvantage was not removed by standardization. Full sex-and-race analyses are reported in Supplementary Materials, Sections 5 and 6.

The benefits of using style-transfer-derived synthetic data also extended to pretrained models. We evaluated BrainAgeNeXt and SynthBA on the synthetic domain test set and compared the results with the real domain predictions discussed in Section 3.2. Paired absolute errors were compared with Wilcoxon signed-rank tests. Synthetic input is out of distribution for BrainAgeNeXt, which was trained on real scans only, and its accuracy declined accordingly: uncorrected *R*^2^ decreased from 0.905 to 0.789 and Beheshti-corrected MAE increased from 4.76 to 5.30 years (*p* = 0.33). Nonetheless, the RFG more than halved, from 1.30 to 0.55 years. SynthBA was itself trained with domain randomization and should in principle already be robust to acquisition variation, yet it still benefited from standardized inputs. Uncorrected *R*^2^ increased from 0.197 to 0.341 for Black subjects and from 0.129 to 0.213 for Asian subjects, while remaining essentially unchanged for White subjects (0.487 to 0.478). Accuracy improved in the same way: overall uncorrected MAE decreased from 14.32 to 13.57 years (*p* = 0.11), but the change was driven primarily by Black subjects (15.96 to 14.12 years, *p* = 0.005), with Asian subjects improving slightly (15.17 to 14.73 years, *p* = 0.82) and White subjects unchanged (11.83 to 11.87 years, *p* = 0.93). The uncorrected RFG narrowed from 4.13 to 2.87 years, and the Beheshti-corrected gap from 1.33 to 0.64 years.

### 3.4 Style-transfer synthesis improves generalization

SuperSynth-derived images are heavily harmonized in intensity and resolution, potentially removing much of the acquisition- and site-specific variation that makes out-of-distribution transfer difficult. This makes them an attractive candidate for improving the generalizability of trained networks. To test this, the all-data model and the fairness-optimized 20% W/60% B/20% A models were evaluated on ADNI3, a cohort previously unseen and differing from the MGH data in acquisition environment, age distribution, and clinical composition; because of these differences, age bias correction was fitted independently within ADNI using CN participants (see Section 7 of Supplementary Materials for details). Calibrated *R*^2^ and Pearson *r* were therefore computed in the CN subgroup, whereas overall MAE was calculated across all clinical groups. Results are summarized in Table 2, with complete correction-stage and clinical-group analyses in Supplementary Materials, Section 7.

**TABLE 2.** External generalization and clinical sensitivity on ADNI3. Overall MAE, CN *R*^2^, and CN Pearson *r* are reported after Beheshti correction. Clinical columns report the fully corrected mean BAG difference from CN and the corresponding MCI effect size. Clinical analyses were not performed for the inadequately calibrated real domain 20% W/60% B/20% A model. MAE and BAG differences are expressed in years.

| Model | Domain | MAE | CN $R^2$ | CN $r$ | SMC-CN BAG | MCI-CN BAG | MCI $d$ |
| --- | --- | --- | --- | --- | --- | --- | --- |
| All data | Real | $4.61 \pm 3.59$ | 0.275 | 0.759 | +0.37 | +2.95 | 0.54 |
| All data | Synthetic | $4.18 \pm 3.40$ | 0.393 | 0.788 | +0.96 | +3.11 | 0.64 |
| 20% W/60% B/20% A | Real | $5.94 \pm 4.51$ | -0.307 | 0.656 | — | — | — |
| 20% W/60% B/20% A | Synthetic | $4.46 \pm 3.23$ | 0.270 | 0.758 | +1.56 | +2.34 | 0.44 |

Note: MCI-CN differences were significant for all three adequately calibrated models ( $p < 0.001$ ); SMC-CN differences were not significant.

In the real domain, the 20% W/60% B/20% A model did not generalize, retaining a negative post linear correction CN *R*^2^ of –0.307 and a Beheshti-corrected MAE of 5.94 *±* 4.51 years. The same configuration trained and evaluated in the synthetic domain recovered performance, with MAE of 4.46 *±* 3.23 years and CN *R*^2^ = 0.270, approaching the all-data synthetic model (MAE 4.18 *±* 3.40 years; CN *R*^2^ = 0.393). The all-data synthetic model also outperformed its real domain counterpart (MAE 4.61 *±* 3.59 years; CN *R*^2^ = 0.275).

Importantly, improved generalization did not come at the cost of clinical signal. The differences in BAG between the CN, SMC and MCI groups were assessed in the three adequately calibrated models (*R*^2^ > 0), excluding the real domain Black-dominant one. In all three, mean BAG increased monotonically from CN to SMC to MCI participants, and MCI participants had significantly higher BAG than CN participants in every model (Welch’s t-test, all *p ≤* 0.001, Cohen’s *d* = 0.44–0.64).

### 3.5 Further mitigation strategies

To further investigate the usefulness of synthetic data, we evaluate its use as a targeted mitigation strategy in the White-dominant configuration, which had the largest real domain fairness gap among the mixed-race fixed-size models. Rather than converting every scan to the synthetic domain, synthetic images were generated only for minority subjects and added to the original real training set. This oversampling approach was compared against race-label injection (Gravina et al. 2026), an established mitigation strategy. Both interventions require access to the training process, so they apply only to models trained here; BrainAgeNeXt and SynthBA were evaluated as released. Architecture, training procedure, validation and test sets, correction pipeline, and fairness metrics were otherwise identical to the primary experiments. Results are summarized in Table 3, with further implementation details in Supplementary Materials, Section 8.

**TABLE 3.** Comparison of bias-mitigation interventions applied to the worst-fairness configuration (60% W/20% B/20% A, Beheshti RFG = 3.00 years). Per-race MAE is shown for all interventions to illustrate redistribution of the prediction gap. Best-composition models included as reference. All MAE values are Beheshti-corrected. Bold values denote the best-performing result within each section, with lower RFG indicating greater fairness.

| Condition | Domain | Overall MAE | RFG | Asian MAE | Black MAE | White MAE |
| --- | --- | --- | --- | --- | --- | --- |
| Baseline: 60% W/20% B/20% A | Real | 7.52 ± 5.08 | 3.00 | 7.96 ± 6.40 | 8.80 ± 6.60 | 5.81 ± 4.23 |
| Pure synthetic: 60% W/20% B/20% A | Synth | **7.09 ± 6.21** | 2.46 | 6.77 ± 6.45 | 8.48 ± 6.97 | 6.02 ± 5.20 |
| Hybrid oversampling† | Mixed | 7.15 ± 5.64 | **1.18** | 7.83 ± 5.78 | 6.97 ± 6.00 | 6.65 ± 5.01 |
| Race-label injection‡ | Real | 7.87 ± 5.35 | 2.22 | 8.97 ± 6.15 | 7.88 ± 5.90 | 6.75 ± 4.80 |
| Reference: 20% W/60% B/20% A | Real | 7.14 ± 5.68 | 0.25 | 7.27 ± 5.62 | 7.13 ± 6.12 | 7.02 ± 5.28 |
| Reference: 20% W/60% B/20% A | Synth | **6.15 ± 5.77** | **0.21** | 6.05 ± 6.07 | 6.14 ± 6.41 | 6.26 ± 4.82 |

†Hybrid: real majority (60% W) supplemented with SynthT1 images of available minority subjects; no new real data collected.

#### Synthetic minority oversampling

The real training set was rebalanced by removing surplus White subjects and adding SuperSynth-derived images of Black and Asian subjects already present in the cohort, until racial balance was reached (*n ≈* 343). This reduced the Beheshticorrected RFG from 3.00 to 1.18 years, a 61% reduction, while also improving overall MAE from 7.52 to 7.15 years. The hybrid model was also fairer than the corresponding fully synthetic White-dominant model, which had an RFG of 2.46 years.

#### Race-label injection

Following Gravina et al. (2026), a one-hot encoding of self-reported race was concatenated to the final pooled feature vector, with all other training settings unchanged. This considerably reduced RFG from 3.00 to 2.22 years at a slight increase in MAE from 7.52 to 7.87 years. Black-participant MAE decreased, whereas Asian-participant MAE increased, shifting rather than uniformly reducing subgroup error.

### 3.6 Interpretability: representational analysis and feature attribution

We examined two complementary aspects of interpretability: how the networks encoded demographic information and which image regions contributed to their predictions. For the first, we tested whether improvements in fairness coincided with changes in the encoding of age and race. Because the network is trained only to predict age, its representations should encode age by construction; race, by contrast, is never supervised, so any race information present must have been picked up directly from the images. Probing both therefore separates what the model was asked to learn from what it learned anyway. We applied t-SNE and linear probing to frozen representations from the best- and worst-fairness configurations in both imaging domains, with additional probes for the all-data and hybrid augmentation models. We extracted features after each convolutional block and from the final embedding layer, and quantified age information using the held-out *R*^2^ of a ridge-regression probe and race information using the balanced accuracy of a multinomial logistic-regression probe. Supplementary Materials, Section 9 provides full methods and t-SNE visualizations.

Across configurations and domains, t-SNE embeddings primarily followed an age gradient, without clear global separation by race or sex. Linear probing nevertheless showed above-chance race decodability from intermediate representations in both domains (permutation *p* < 0.05; Figure 4). Race-decoding accuracy remained broadly similar between models with markedly different fairness gaps, including the best- and worst-fairness configurations. Age decodability followed a different pattern: synthetic domain representations generally showed higher age *R*^2^ than real domain representations across most network depths, although this difference narrowed in the deepest layers.

**FIGURE 4.**
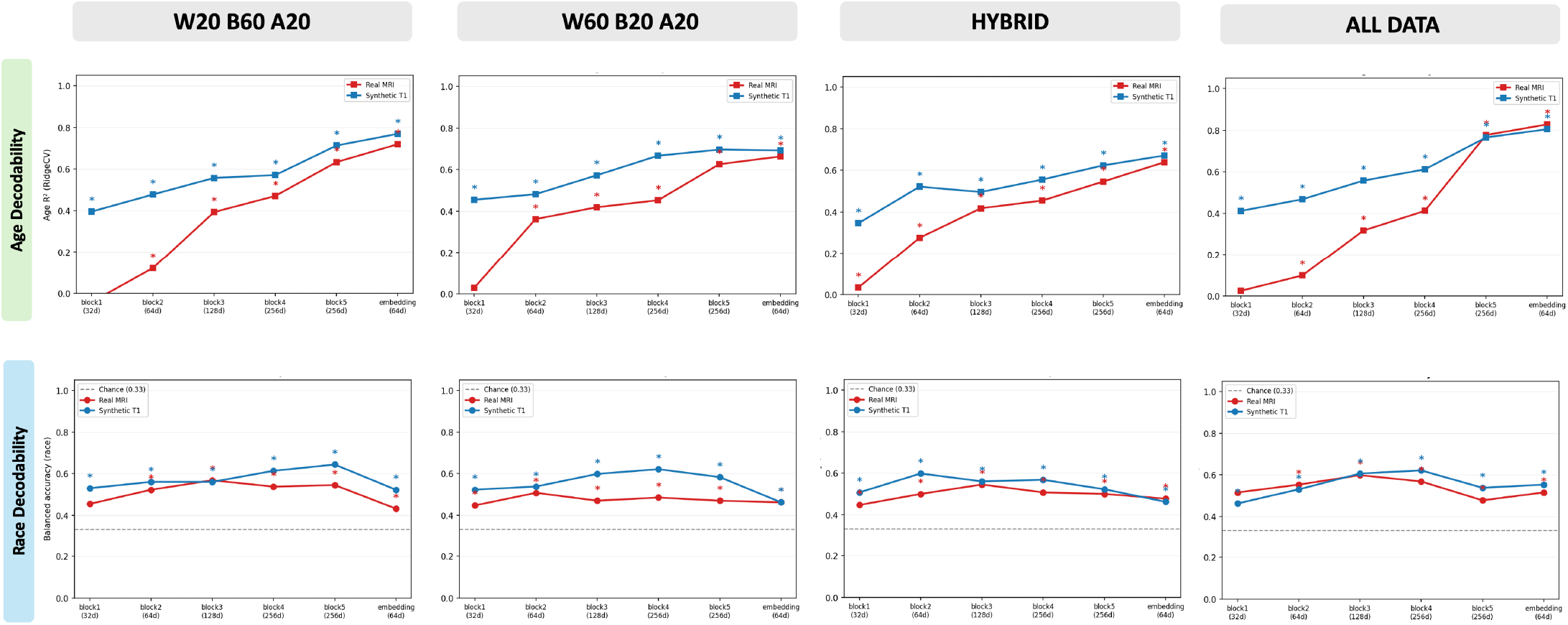
Linear probing of race and age information across SFCN network depth for four training configurations. Columns show the Black-dominant 20% W/60% B/20% A model, the White-dominant 60% W/20% B/20% A model, the hybrid model, and the all-data model, respectively. Measurements are taken at each block of the SFCN. **Upper row:** Age decodability, measured as the held-out test *R*^2^ of a ridge-regression probe trained on validation set representations. **Lower row:** Race decodability, measured as the held-out test balanced accuracy of a multinomial logistic-regression probe trained on validation set representations; the horizontal dashed line denotes chance performance for the three-class task (1/3). In both rows, results are shown separately for real MRI and synthetic T1 representations, and asterisks indicate decoding above the permutation null (*p* < 0.05, 1,000 label shuffles).

To complement these representational analyses, we used Integrated Gradients (Sundararajan et al. 2017) to identify spatial features contributing to brain age predictions and determine whether synthetic standardization altered their use. We compared the Black-dominant and White-dominant models, which showed the most contrasting fairness outcomes among fixed-size configurations, across three contrasts: race (Black minus White), domain (synthetic minus real in Black subjects), and age (18–37 minus 78–90 years). We summarized attribution differences using the SuperSynth segmentation of the MNI template across intracranial and extracerebral ROIs.

Domain differences showed greater consistency across models than racial differences. Synthetic images produced stronger contributions from the caudate and ventricular system, including the fourth ventricle (+1.22 *×* 10^−4^ in the Black-dominant and +1.65 *×* 10^−4^ in the White-dominant model), whereas real images showed stronger contributions from cerebellar white matter and the accumbens area. Age produced the largest attribution differences. In real images, the bilateral caudate and fornix contributed more strongly for older subjects in both models, with caudate differences of approximately –1.0 to –1.4 *×* 10^−4^. Synthetic images also concentrated age-related differences in the fornix, although less consistently across models. Racial differences remained smaller, more localized, and less consistent across models and domains.

When extracerebral ROIs were included, some of the largest signed attribution differences were observed in vascular, ocular, muscular, and skeletal tissues. Domain-related effects were the most consistent: the arterial ROI showed stronger attribution in synthetic images in both configurations (+7.59 *×* 10^−4^ in the Black-dominant model and +5.41 *×* 10^−4^ in the White-dominant model). Age-related differences were also observed in cancellous bone, optic nerves, ocular structures, and extraocular muscles. In contrast, racial and age-related arterial differences varied in direction across imaging domains and training compositions. Section 10 of Supplementary Materials provides the full methods and ROI-level results.

## 4 DISCUSSION

Prior work has largely framed demographic disparities in brain age prediction as a consequence of training set imbalance (Piçarra and Glocker 2023, Dempsey et al. 2023, Adkins and Hanson 2025, Aguzin Parrilli and Belzunce 2026). Separately, appearance harmonization has been developed to reduce scanner- and site-related variation, motivated primarily by accuracy and robustness (Dewey et al. 2019, Zuo et al. 2021, Iglesias et al. 2023). Our findings connect these two lines of work by showing that standardization can also function as a fairness intervention. Because our paired real-synthetic design holds participant anatomy and demographic composition constant, and because the domain effect holds also in models trained on a single racial group, the fairness improvement is attributable directly to the domain change.

A plausible explanation is that style-transfer standardization reduces reliance on acquisition-related shortcuts (Geirhos et al. 2020, DeGrave et al. 2021). Scanner, protocol, site, and other acquisition characteristics are most often unevenly distributed across different demographic groups and can therefore become shortcuts for the model to assign subgroup membership. The representational analyses suggest that improved fairness does not arise simply from removing demographic information: race remained decodable after appearance harmonization. In any case, IG attribution maps show that the relative importance of regions contributing to age prediction shifted between the two domains, with age-relevant ROIs preserved in both, highlighting regions in line with prior brain age attribution work (Sundararajan et al. 2017, Guo et al. 2024, Hofmann et al. 2025). Notably, the largest domain differences appeared outside the brain, in vascular and other extracerebral tissue, and were consistent in direction across both configurations, whereas racial and age-related differences in the same regions varied in sign. These tissues are plausible contributors to age prediction, since arteries, bone, orbital structures and extraocular muscles all change measurably with age (Cali et al. 2023, Keenan et al. 2009, Watanabe et al. 2017, Paskhover et al. 2017). Taken together, these results are more consistent with standardization changing what information the model relies on than with it nulling the correlation between features and race.

Another insight comes from the SynthBA results. Despite being trained entirely on images with randomized intensities, which should discourage reliance on acquisition-specific appearance, SynthBA showed poor racial fairness on real images but improved when evaluated on synthetic scans. This suggests that robustness to appearance variation does not necessarily translate into fairness. Domain-randomized training encourages the model to become robust to differences in image contrast and resolution, whereas appearance harmonization removes these differences before prediction. The improvement on synthetic scans therefore indicates that appearance-related factors may continue to contribute to racial disparities even when models are trained to be robust to them.

Importantly, ours is not a universal fairness solution. Training composition remained influential, the reduction was specific to racial rather than sex disparities, and intersectional disparities persisted, with Black Female subjects showing the highest errors across the synthetic domain configurations. These findings suggest that acquisition-related variation explains only one component of demographic disparity and that different protected attributes may reflect different mechanisms.

From a practical perspective, our results motivate interventions that act on the imaging pipeline rather than treating demographic labels themselves as corrective inputs. This distinction is also conceptually important because race is a socially constructed and context-dependent category rather than a categorical variable. Self-reported identities may be multiracial, may not map cleanly onto mutually exclusive categories, and may be recorded inconsistently across clinical systems. Using race directly as a model input therefore requires reducing complex identities to fixed categories and assumes that those labels are both available and meaningful at deployment.

Style-transfer standardization avoids this requirement and can be applied retrospectively to already collected scans using open-source tools (Iglesias et al. 2023, Billot et al. 2023a). Synthetic minority oversampling provides an additional option when training data are imbalanced (Jackson et al. 2025). Demographic information is still required during dataset construction to identify underrepresented groups, but unlike race-label injection (Gravina et al. 2026), race is not encoded as a predictive feature and is not required at deployment. Taken together, these results support style-transfer standardization as a promising approach to reducing demographic disparities and motivate further investigation into the mechanisms and broader implications of this effect.

A limitation shared by all brain age models is that equal prediction error does not necessarily imply equal biological aging. Differences in brain aging may arise from education, socioeconomic conditions, vascular health, environmental exposures, chronic stress, healthcare access, and other lived experiences (Farina et al. 2023, Levine and Crimmins 2014). A residual group difference could therefore reflect either undesirable acquisition or model bias, or genuine differences in brain aging associated with socially patterned health. Consequently, group differences in prediction error cannot be attributed unambiguously to algorithmic unfairness.

Within this context, the balanced test set of 44 subjects per racial group supported paired configuration-level tests of the domain effect, although within-model subgroup analyses remain exploratory. The cohort was also limited to participants who self-identified as Asian, Black, or White. Future studies using larger and more diverse datasets with richer metadata could enable subject-level inference, extend evaluation to additional demographic groups, and better account for potential confounding factors.

At the same time, the study examined thirteen training compositions and two imaging domains using a widely reproduced architecture and a well-characterized synthesis framework, and external benchmarks suggest that the observed domain effect is not specific to our network. Whether the magnitude of this effect depends on model architecture remains an open question, as do diffusion- and GAN-based alternatives for synthesis.

Finally, the study focused on cross-sectional prediction error, a well-established proxy for the clinical utility of BAG. Direct evaluation of whether BAG provides comparable prognostic and disease-related information across groups will require longitudinal outcome data and remains an important direction for future work.

## 5 CONCLUSION

In this article, we have shown that synthetic scans mimicking a 1 mm T1-weighted acquisition can reduce racial disparities in brain age prediction while improving accuracy and robustness. Across a wide range of training compositions, models trained and evaluated in the synthetic domain consistently exhibited smaller racial performance gaps, better overall accuracy, and improved out-of-distribution generalization compared with their real-domain counterparts. Similar improvements in external pretrained models suggest that these benefits are not specific to a particular training strategy or architecture. Representational and attribution analyses further indicated that style-transfer standardization alters the features used for age prediction rather than simply removing race-associated information. Finally, augmenting an imbalanced training set with synthetic images of minority subjects already present in the cohort closed 61% of the racial performance gap, highlighting a practical strategy for improving equity without recruiting additional participants.

The software used for 1 mm T1-weighted synthesis (SuperSynth) is openly available through FreeSurfer and can be applied retrospectively to scans already acquired. The interventions proposed here therefore require no additional subjects and, once a model has been trained,no knowledge of race at deployment. By reducing acquisition-related variability while preserving anatomical information, style-transfer harmonization holds promise to facilitate not only more robust and reproducible neuroimaging applications, but also more equitable ones.

## AUTHOR CONTRIBUTIONS

Conceptualization: V.B., J.E.I. Data curation: D.Z. Formal analysis: V.B. Funding acquisition: J.E.I. Project administration: J.E.I. Software: V.B., D.Z. Supervision: D.Z., J.E.I. Visualization: V.B. Writing—original draft: V.B. Writing—review and editing: all authors.

## ACKNOWLEDGMENTS

This work was primarily supported by NIH grant 1RF1AG080371. Additional support was received from NIH grants 1R01AG070988, 1UM1MH130981, 1R21NS138995, U19AG060909, and from the MIT-MGB Seed Program sponsored by MIT HEALS. Data collection and sharing for the Alzheimer’s Disease Neuroimaging Initiative (ADNI) is funded by the National Institute on Aging (National Institutes of Health Grant U19 AG024904). The grantee organization is the Northern California Institute for Research and Education. In the past, ADNI has also received funding from the National Institute of Biomedical Imaging and Bioengineering, the Canadian Institutes of Health Research, and private sector contributions through the Foundation for the National Institutes of Health (FNIH) including generous contributions from the following: AbbVie, Alzheimer’s Association; Alzheimer’s Drug Discovery Foundation; Araclon Biotech; BioClinica, Inc.; Biogen; Bristol-Myers Squibb Company; CereSpir, Inc.; Cogstate; Eisai Inc.; Elan Pharmaceuticals, Inc.; Eli Lilly and Company; EuroImmun; F. Hoffmann-La Roche Ltd and its affiliated company Genentech, Inc.; Fujirebio; GE Healthcare; IXICO Ltd.; Janssen Alzheimer Immunotherapy Research & Development, LLC.; Johnson & Johnson Pharmaceutical Research &Development LLC.; Lumosity; Lundbeck; Merck & Co., Inc.; Meso Scale Diagnostics, LLC.; NeuroRx Research; Neurotrack Technologies; Novartis Pharmaceuticals Corporation; Pfizer Inc.; Piramal Imaging; Servier; Takeda Pharmaceutical Company; and Transition Therapeutics.

## CONFLICT OF INTEREST

The authors declare no potential conflict of interests.

## Notes

### Competing Interest Statement

The authors have declared no competing interest.

## REFERENCES

Adkins, D.J. & Hanson, J.L. (2025) Racial and ethnic disparities in brain age algorithm performance: Investigating bias across six popular methods. medRxiv,. 10.1101/2025.09.18.25336117.

Adleberg, J., Wardeh, A., Doo, F.X., Marinelli, B., Cook, T.S., Mendelson, D.S. et al. (2022) Predicting patient demographics from chest radiographs with deep learning. Journal of the American College of Radiology, 19(10), 1151–1161. 10.1016/j.jacr.2022.06.008.

Aguzin Parrilli, L.J. & Belzunce, M.A. (2026) Bias and generalizability of brain age prediction models: A multi-cohort evaluation with anatomical and interpretability insights. Imaging Neuroscience, 4, IMAG.a.1164. 10.1162/IMAG.a.1164.

Alain, G. & Bengio, Y. (2016) Understanding intermediate layers using linear classifier probes.

Alqarni, M., Jones, E.L., Ribeiro, L., Verma, H., Cooper, S., Mullassery, V. et al. (2025) An investigation of race bias in deep learning-based segmentation of prostate MRI images. Scientific Reports, 15(1), 43427. 10.1038/s41598-025-26189-5.

Banerjee, I., Bhattacharjee, K., Burns, J.L., Trivedi, H., Purkayastha, S., Seyyed-Kalantari, L. et al. (2023) Shortcuts causing bias in radiology artificial intelligence: Causes, evaluation, and mitigation. Journal of the American College of Radiology : JACR, 20(9), 842–851. 10.1016/j.jacr.2023.06.025.

Beheshti, I., Nugent, S., Potvin, O. & Duchesne, S. (2019) Bias-adjustment in neuroimaging-based brain age frameworks: A robust scheme. NeuroImage: Clinical, 24, 102063. 10.1016/j.nicl.2019.102063.

Benjamini, Y. & Hochberg, Y. (1995) Controlling the false discovery rate: A practical and powerful approach to multiple testing. Journal of the Royal Statistical Society: Series B, 57(1), 289–300. 10.1111/j.2517-6161.1995.tb02031.x.

Billot, B., Greve, D.N., Puonti, O., Thielscher, A., Van Leemput, K., Fischl, B. et al. (2023) SynthSeg: Segmentation of brain MRI scans of any contrast and resolution without retraining. Medical Image Analysis, 86, 102789. 10.1016/j.media.2023.102789.

Billot, B., Magdamo, C., Cheng, Y., Arnold, S.E., Das, S. & Iglesias, J.E. (2023) Robust machine learning segmentation for large-scale analysis of heterogeneous clinical brain MRI datasets. Proceedings of the National Academy of Sciences, 120(9), e2216399120. 10.1073/pnas.2216399120.

Biondo, F., Bennallick, C., Martin, S.A., Puglisi, L., Booth, T.C., Wood, D.A. et al. (2025) Brain-age in ultra-low-field MRI: how well does it work? medRxiv, 2025–10. 10.1101/2025.10.19.25338298.

Brown, A., Tomasev, N., Freyberg, J. et al. (2023) Detecting shortcut learning for fair medical AI using shortcut testing. Nature Communications, 14, 4314. 10.1038/s41467-023-39902-7.

Cali, R.J., Bhatt, R.R., Thomopoulos, S.I., Gadewar, S., Ba Gari, I., Chattopadhyay, T. et al. (2023) The influence of brain MRI defacing algorithms on brain-age predictions via 3D convolutional neural networks. bioRxiv, 2023.04.28.538724. 10.1101/2023.04.28.538724, preprint.

Cole, J.H. & Franke, K. (2017) Predicting age using neuroimaging: Innovative brain ageing biomarkers. Trends in Neurosciences, 40(12), 681–690. 10.1016/j.tins.2017.10.001.

Cole, J.H., Ritchie, S.J., Bastin, M.E., Valdés Hernández, M.C., Muñoz Maniega, S., Royle, N. et al. (2018) Brain age predicts mortality. Molecular Psychiatry, 23(5), 1385–1392. 10.1038/mp.2017.62.

Danaee, G., Niethammer, M., Rushmore, J. & Bouix, S. (2025) Investigating demographic bias in brain MRI segmentation: A comparative study of deep-learning and non-deep-learning methods. Machine Learning for Biomedical Imaging, 3, 792–807. 10.59275/j.melba.2025-d1g3.

de Lange, A.M.G., Antürk, M., Rokicki, J., Han, L.K.M., Franke, K., Alnaes, D. et al. (2022) Mind the gap: Performance metric evaluation in brain-age prediction. Human Brain Mapping, 43(10), 3113–3129. 10.1002/hbm.25837.

DeGrave, A.J., Janizek, J.D. & Lee, S.I. (2021) AI for radiographic COVID-19 detection selects shortcuts over signal. Nature Machine Intelligence, 3(7), 610–619. 10.1038/s42256-021-00338-7.

Dempsey, D.A., Deardorff, R., Wu, Y.C., Yu, M., Apostolova, L.G., Brosch, J. et al. (2023) BrainAGE estimation: Influence of field strength, voxel size, race, and ethnicity. medRxiv, 2023.12.05.23299222. 10.1101/2023.12.05.23299222.

Dewey, B.E., Zhao, C., Reinhold, J.C. et al. (2019) Deepharmony: A deep learning approach to contrast harmonization across scanner changes. Magnetic Resonance Imaging, 64, 160–170. 10.1016/j.mri.2019.05.041.

Dörfel, R.P., Arenas-Gomez, J.M., Fisher, P.M., Ganz, M., Knudsen, G.M., Svensson, J.E. et al. (2023) Prediction of brain age using structural magnetic resonance imaging: A comparison of accuracy and test-retest reliability of publicly available software packages. Human Brain Mapping, 44(17), 6139–6148. 10.1002/hbm.26502.

Du, Y., Xue, Y., Dharmakumar, R. & Tsaftaris, S.A. (2023) Unveiling fairness biases in deep learning-based brain MRI reconstruction. arXiv, (arXiv:2309.14392). 10.1007/978-3031-45249-9_10.

Duffy, G., Clarke, S.L., Christensen, M., He, B., Yuan, N., Cheng, S. et al. (2022) Confounders mediate AI prediction of demographics in medical imaging. npj Digital Medicine, 5, 188. 10.1038/s41746-022-00720-8.

Farina, M.P., Kim, J.K. & Crimmins, E.M. (2023) Racial/ethnic differences in biological aging and their life course socioeconomic determinants: the 2016 health and retirement study. Journal of aging and health, 35(3), 209–220. 10.1177/08982643221120743.

Fischl, B. (2012) Freesurfer. NeuroImage, 62(2), 774–781. 10.1016/j.neuroimage.2012.01.021.

Franke, K., Ziegler, G., Klöppel, S. & Gaser, C. (2010) Estimating the age of healthy subjects from T1-weighted MRI scans using kernel methods: exploring the influence of various parameters. NeuroImage, 50(3), 883–892. 10.1016/j.neuroimage.2010.01.005.

Geirhos, R., Jacobsen, J.H., Michaelis, C., Zemel, R., Brendel, W., Bethge, M. et al. (2020) Shortcut learning in deep neural networks. Nature Machine Intelligence,. 10.1038/s42256-020-00257-z.

Gichoya, J.W., Banerjee, I., Bhimireddy, A.R., Burns, J.L., Celi, L.A., Chen, L.C. et al. (2022) AI recognition of patient race in medical imaging: a modelling study. The Lancet Digital Health, 4(6), e406–e414. 10.1016/S2589-7500(22)00063-2.

Gravina, M., Pontillo, G., Shawa, Z., Cole, J.H. & Sansone, C. (2026) Assessing demographic bias in brain age prediction models using multiple deep learning paradigms. Pattern Recognition Letters, 199, 246–253. 10.1016/j.patrec.2025.11.029.

Guo, K.H., Chaudhari, N.N., Jafar, T., Chowdhury, N.F., Bogdan, P., Irimia, A. et al. (2024) Anatomic interpretability in neuroimage deep learning: Saliency approaches for typical aging and traumatic brain injury. Neuroinformatics, 22, 591–606. 10.1007/s12021-024-09694-2.

Hoffmann, M. (2025) Domain-randomized deep learning for neuroimage analysis: Selecting training strategies, navigating challenges, and maximizing benefits. IEEE Signal Processing Magazine, 42(4), 78–90. 10.1109/MSP.2025.3590806.

Hofmann, S.M., Goltermann, O., Scherf, N., Müller, K.R., Löffler, M., Villringer, A. et al. (2025) The utility of explainable AI for MRI analysis: Relating model predictions to neuroimaging features of the aging brain. Imaging Neuroscience, 3, imag_a_00497. 10.1162/imag_a_00497.

Iglesias, J.E., Billot, B., Balbastre, Y., Magdamo, C., Arnold, S.E., Das, S. et al. (2023) SynthSR: A public AI tool to turn heterogeneous clinical brain scans into high-resolution T1-weighted images for 3d morphometry. Science Advances, 9(5), eadd3607. 10.1126/sciadv.add3607.

Ioannou, S., Chockler, H., Hammers, A. & King, A.P. (2022) A study of demographic bias in CNN-based brain MR segmentation. Machine Learning in Clinical Neuroimaging, 13–22. 10.1007/978-3-031-17899-3_2.

Ioffe, S. & Szegedy, C. Batch normalization: Accelerating deep network training by reducing internal covariate shift. In: Proceedings of the 32nd International Conference on Machine Learning (ICML). Vol. 37, 2015, pp. 448–456.

Jack, C.R.J., Bernstein, M.A., Fox, N.C., Thompson, P., Alexander, G., Harvey, D. et al. (2008) The alzheimer’s disease neuroimaging initiative (adni): MRI methods. Journal of Magnetic Resonance Imaging, 27(4), 685–691. 10.1002/jmri.21049.

Jackson, N.J., Yan, C. & Malin, B.A. (2025) Enhancement of fairness in AI for chest x-ray classification. AMIA Annual Symposium Proceedings, 2024, 551–560.

Keenan, N.G., Locca, D., Varghese, A., Roughton, M., Gatehouse, P.D., Hooper, J. et al. (2009) Magnetic resonance of carotid artery ageing in healthy subjects. Atherosclerosis, 205(1), 168–173. 10.1016/j.atherosclerosis.2008.11.018.

Kruskal, W.H. & Wallis, W.A. (1952) Use of ranks in one-criterion variance analysis. Journal of the American Statistical Association, 47(260), 583–621. 10.1080/01621459.1952.10483441.

La Rosa, F., Dos Santos Silva, J., Dereskewicz, E., Invernizzi, A., Cahan, N., Galasso, J. et al. (2025) BrainAgeNeXt: Advancing brain age modeling for individuals with multiple sclerosis. Imaging Neuroscience, 3, imag_a_00487. 10.1101/2024.08.10.24311686.

Levine, M. & Crimmins, E. (2014) Evidence of accelerated aging among african americans and its implications for mortality. Social Science Medicine, 118, 27–32. 10.1016/j.socscimed.2014.07.022.

Liang, H., Zhang, F. & Niu, X. (2019) Investigating systematic bias in brain age estimation with application to post-traumatic stress disorders. Human Brain Mapping, 40(11), 3143–3152. 10.1002/hbm.24588.

Litjens, G., Kooi, T., Bejnordi, B.E., Setio, A.A.A., Ciompi, F., Ghafoorian, M. et al. (2017) A survey on deep learning in medical image analysis, 42, 60–88. 10.1016/j.media.2017.07.005.

Liu, M., Zhu, A.H., Maiti, P., Thomopoulos, S.I., Gadewar, S., Chai, Y. et al. (2023) Style transfer generative adversarial networks to harmonize multisite MRI to a single reference image to avoid overcorrection. Human Brain Mapping, 44(14), 4875–4892. 10.1002/hbm.26422.

Liu, P., Zemlyanker, D., Gopinath, K., Cheng, Y., He, Y., Izquierdo-Garcia, D. et al. (2025) The normalizing properties of intracranial volume across race and sex. Brain Communications, 7(4), fcaf271. 10.1093/braincomms/fcaf271.

Paskhover, B., Durand, D., Kamen, E. & Gordon, N.A. (2017) Patterns of change in facial skeletal aging. JAMA Facial Plastic Surgery, 19(5), 413–417. 10.1001/jamafacial.2017.0743.

Pedregosa, F., Varoquaux, G., Gramfort, A. et al. (2011) Scikit-learn: Machine learning in Python. Journal of Machine Learning Research, 12, 2825–2830.

Peng, H., Gong, W., Beckmann, C.F., Vedaldi, A. & Smith, S.M. (2021) Accurate brain age prediction with lightweight deep neural networks. Medical Image Analysis, 68, 101871. 10.1016/j.media.2020.101871.

Piçarra, C. & Glocker, B. Analysing race and sex bias in brain age prediction. In: Clinical Image-Based Procedures, Fairness of AI in Medical Imaging, and Ethical and Philosophical Issues in Medical Imaging. Vol. 14242 of Lecture Notes in Computer Science, 2023. Cham: Springer.

Puglisi, L., Rondinella, A., Meo, L.D., Guarnera, F., Battiato, S. & Ravì, D. (2024) SynthBA: Reliable brain age estimation across multiple MRI sequences and resolutions. 2024 IEEE International Conference on Metrology for eXtended Reality, Artificial Intelligence and Neural Engineering (MetroXRAINE), 559–564. 10.1109/MetroXRAINE62247.2024.10796114.

Puyol-Antón, E., Ruijsink, B., Mariscal Harana, J., Piechnik, S.K., Neubauer, S., Petersen, S.E. et al. (2022) Fairness in cardiac magnetic resonance imaging: Assessing sex and racial bias in deep learning-based segmentation. 9, 859310. 10.3389/fcvm.2022.859310.

Ricci Lara, M.A., Echeveste, R. & Ferrante, E. (2022) Addressing fairness in artificial intelligence for medical imaging. Nature Communications, 13(1), 4581. 10.1038/s41467-022-32186-3.

Seyyed-Kalantari, L., Zhang, H., McDermott, M.B.A., Chen, I.Y. & Ghassemi, M. (2021) Underdiagnosis bias of artificial intelligence algorithms applied to chest radiographs in under-served patient populations. Nature Medicine, 27(12), 2176–2182.

Shen, D., Wu, G. & Suk, H.I. (2017) Deep learning in medical image analysis. Annual Review of Biomedical Engineering, 19, 221–248. 10.1146/annurev-bioeng-071516-044442.

Smith, S.M., Vidaurre, D., Alfaro-Almagro, F., Nichols, T.E. & Miller, K.L. (2019) Estimation of brain age delta from brain imaging. NeuroImage, 200, 528–539. 10.1016/j.neuroimage.2019.06.017.

Sundararajan, M., Taly, A. & Yan, Q. (2017) Axiomatic attribution for deep networks. https://arxiv.org/abs/1703.01365

Treder, M.S., Shock, J.P., Stein, D.J., du Plessis, S., Seedat, S. & Tsvetanov, K.A. (2021) Correlation constraints for regression models: Controlling bias in brain age prediction. Frontiers in Psychiatry, Volume 12 - 2021. 10.3389/fpsyt.2021.615754.

Valdes-Hernandez, P.A., Laffitte Nodarse, C., Peraza, J.A., Cole, J.H. & Cruz-Almeida, Y. (2023) Toward MR protocol-agnostic, unbiased brain age predicted from clinical-grade MRIs. Scientific Reports, 13(1), 19570. 10.1038/s41598-023-47021-y.

van Breugel, B., Kyono, T., Berrevoets, J. & van der Schaar, M. (2021) Decaf: Generating fair synthetic data using causally-aware generative networks. https://arxiv.org/abs/2110.12884

van Breugel, B., Liu, T., Oglic, D. & van der Schaar, M. (2024) Synthetic data in biomedicine via generative artificial intelligence. Nature Reviews Bioengineering, 2, 991–1004. 10.1038/s44222-024-00245-7.

van Breugel, B., Qian, Z. & Schaar, M.V.D. (2023) Synthetic data, real errors: How (not) to publish and use synthetic data. Proceedings of the 40th International Conference on Machine Learning, 34793–34808.

van Breugel, B., Seedat, N., Imrie, F. & van der Schaar, M. (2023) Can you rely on your model evaluation? improving model evaluation with synthetic test data. https://arxiv.org/abs/2310.16524

van der Maaten, L. & Hinton, G. (2008) Visualizing data using t-SNE. Journal of Machine Learning Research, 9(86), 2579–2605. http://jmlr.org/papers/v9/vandermaaten08a.html

Virtanen, P., Gommers, R., Oliphant, T.E. et al. (2020) SciPy 1.0: Fundamental algorithms for scientific computing in Python. Nature Methods, 17, 261–272. 10.1038/s41592-019-0686-2.

Wang, R., Kuo, P.C., Chen, L.C., Seastedt, K.P., Gichoya, J.W. & Celi, L.A. (2024) Drop the shortcuts: image augmentation improves fairness and decreases AI detection of race and other demographics from medical images. EBioMedicine, 102. 10.1016/j.ebiom.2024.105047.

Watanabe, M., Buch, K., Fujita, A., Jara, H., Qureshi, M.M. & Sakai, O. (2017) Quantitative MR imaging of intraorbital structures: Tissue-specific measurements and age dependency compared to extra-orbital structures using multispectral quantitative MR imaging. Orbit, 36(4), 189–196. 10.1080/01676830.2017.1310254.

Weiner, M.W., Veitch, D.P., Aisen, P.S., Beckett, L.A., Cairns, N.J., Green, R.C. et al. (2017) The alzheimer’s disease neuroimaging initiative 3: Continued innovation for clinical trial improvement. Alzheimer’s & Dementia, 13(5), 561–571. 10.1016/j.jalz.2016.10.006, epub 2016 Dec 5.

Wilcoxon, F. (1945) Individual comparisons by ranking methods. Biometrics Bulletin, 1(6), 80–83. 10.2307/3001968.

Wu, Y. & He, K. Group normalization. In: Computer Vision – ECCV 2018, 2018. : Springer.

Xu, Z., Li, J., Yao, Q., Li, H., Zhao, M. & Zhou, S.K. (2024) Addressing fairness issues in deep learning-based medical image analysis: a systematic review. npj Digital Medicine, 7(1), 286. 10.1038/s41746-024-01276-5.

Yang, Y., Zhang, H., Gichoya, J.W. et al. (2024) The limits of fair medical imaging AI in real-world generalization. Nature Medicine, 30, 2838–2848. 10.1038/s41591-024-03113-4.

Zhang, B., Zhang, S., Feng, J. & Zhang, S. (2023) Age-level bias correction in brain age prediction. NeuroImage : Clinical, 37, 103319. 10.1016/j.nicl.2023.103319.

Zhang, L. & Wu, X. (2017) Anti-discrimination learning: a causal modeling-based framework. International Journal of Data Science and Analytics, 4, 1–16. 10.1007/s41060-017-0058-x.

Zhang, R., Yi, F., Mao, H., Huang, Z., Wang, K. & Zhang, J. (2025) Brain age gap as a predictive biomarker that links aging, lifestyle, and neuropsychiatric health. Communications Medicine, 5(1), 441. 10.1038/s43856-025-01100-5.

Zuo, L., Dewey, B.E., Liu, Y. et al. (2021) Unsupervised MR harmonization by learning disentangled representations using information bottleneck theory. NeuroImage, 243, 118569. 10.1016/j.neuroimage.2021.118569.

