## Supplementary Material for "Improving racial fairness in brain age models using style-transfer synthesis"

#### Contents


S1 BatchNorm versus GroupNorm stability comparison 2

S2 Age-bias correction and BAG–age correlations 6

S3 Performance across all training configurations 8

S4 External benchmark models 10

S5 Statistical significance of racial and sex fairness gaps 11

S6 Race × sex subgroup performance 13

S7 External validation on ADNI3 19

S8 Race-label injection implementation 20

S9 Representational analysis 21

S10 Integrated gradients 24

### S1 BatchNorm versus GroupNorm stability comparison

#### Methods

To justify the use of GroupNorm over BatchNorm3D (Ioffe and Szegedy, 2015) throughout the composition sweep, two fixed-size sweep configurations ($n \sim 343$), 100% Asian and 60% W/20% B/20% A, were each trained with both normalization schemes, holding all other architectural and training hyperparameters constant. Five independent seeds were trained per normalization choice and configuration, in the real domain. Seed-to-seed training stability was quantified as the coefficient of variation (CV) of per-epoch validation loss, computed after excluding the Lightning sanity-check epoch. Final test performance was assessed on uncorrected predictions (MAE, $R^2$) for each of the five seeds.

The number of groups was fixed at $G = 8$ throughout. $G$ must divide the channel count at every layer, which for this architecture (32, 64, 128, 256, 256, and 64 channels) admits only $G \in \{1, 2, 4, 8, 16, 32\}$. $G = 8$ yields 4 to 32 channels per group across the network, avoiding both degenerate limits ($G = 1$ reduces to LayerNorm and $G = C$ to InstanceNorm, the two settings at which Wu and He (2018) report degraded performance) and falls within the range they recommend, since they found accuracy to be otherwise insensitive to $G$.

#### Results

At $n = 343$, BatchNorm's validation-loss CV exceeded GroupNorm's for both configurations tested (100% Asian: 63.8% vs 45.1%; 60% W/20% B/20% A: 71.7% vs 44.4%). Early-stopping epoch was also more variable across seeds under BatchNorm for the 60% W/20% B/20% A configuration specifically (range 19–61 vs 60–88 epochs for GroupNorm), though this pattern did not hold as clearly for the 100% Asian configuration, indicating that composition-specific factors beyond sample size alone likely contribute to BatchNorm's instability. Per-seed loss curves are shown in Figure S1 (BatchNorm) and Figure S2 (GroupNorm).

This training-time instability translated directly into worse and less reliable final test performance (Table S1). Averaged across five seeds, BatchNorm's uncorrected test MAE exceeded GroupNorm's by 2.3–4.3 years (100% Asian: 12.64 vs 9.54 years; 60% W/20% B/20% A: 14.76 vs 10.48 years), with 3–7× higher seed-to-seed standard deviation (100% Asian: $\pm 3.57$ vs $\pm 0.53$ years; 60% W/20% B/20% A: $\pm 1.74$ vs $\pm 0.58$ years). Uncorrected $R^2$ showed the same pattern (100% Asian: $0.394 \pm 0.350$ vs $0.662 \pm 0.029$; 60% W/20% B/20% A: $0.223 \pm 0.195$ vs $0.604 \pm 0.043$), and one BatchNorm seed (100% Asian, seed 5) produced a negative $R^2$ ($-0.286$), performing worse than predicting the sample mean age for every subject, indicating a training failure.

These results confirm that at the $n \sim 343$ scale used throughout the composition sweep, BatchNorm's instability is not limited to noisier training dynamics that average out by test time: it produces materially worse and substantially less reliable final models. This provides empirical justification for the use of GroupNorm as the default normalization choice throughout the composition sweep, consistent with GroupNorm's known robustness to small and compositionally variable mini-batches (Wu and He, 2018).

**Table S1:** Per-seed uncorrected test performance, BatchNorm vs GroupNorm, at $n = 343$. All metrics computed on the balanced test set ($n = 132$ subjects) prior to any age-bias correction.

| Configuration | Norm | Seed | MAE | $R^2$ | RMSE |
|---|---|---|---|---|---|
| 100% Asian | BatchNorm | 1 | 11.46 | 0.534 | 14.17 |
| | | 2 | 12.77 | 0.424 | 15.77 |
| | | 3 | 9.72 | 0.657 | 12.17 |
| | | 4 | 9.83 | 0.641 | 12.44 |
| | | 5 | 19.43 | −0.286 | 23.55 |
| | *mean ± std* | | **12.64 ± 3.57** | **0.394 ± 0.350** | 15.62 ± 4.31 |
| 100% Asian | GroupNorm | 1 | 9.44 | 0.677 | 11.80 |
| | | 2 | 9.65 | 0.661 | 12.10 |
| | | 3 | 9.02 | 0.687 | 11.62 |
| | | 4 | 10.50 | 0.606 | 13.04 |
| | | 5 | 9.11 | 0.679 | 11.77 |
| | *mean ± std* | | **9.54 ± 0.53** | **0.662 ± 0.029** | 12.07 ± 0.55 |
| 60% W/20% B/20% A | BatchNorm | 1 | 14.55 | 0.273 | 17.71 |
| | | 2 | 12.77 | 0.430 | 15.68 |
| | | 3 | 13.10 | 0.413 | 15.91 |
| | | 4 | 17.37 | −0.065 | 21.43 |
| | | 5 | 16.04 | 0.065 | 20.08 |
| | *mean ± std* | | **14.76 ± 1.74** | **0.223 ± 0.195** | 18.16 ± 2.31 |
| 60% W/20% B/20% A | GroupNorm | 1 | 10.41 | 0.602 | 13.10 |
| | | 2 | 11.11 | 0.570 | 13.62 |
| | | 3 | 10.80 | 0.578 | 13.49 |
| | | 4 | 9.40 | 0.688 | 11.61 |
| | | 5 | 10.67 | 0.583 | 13.41 |
| | *mean ± std* | | **10.48 ± 0.58** | **0.604 ± 0.043** | 13.05 ± 0.70 |

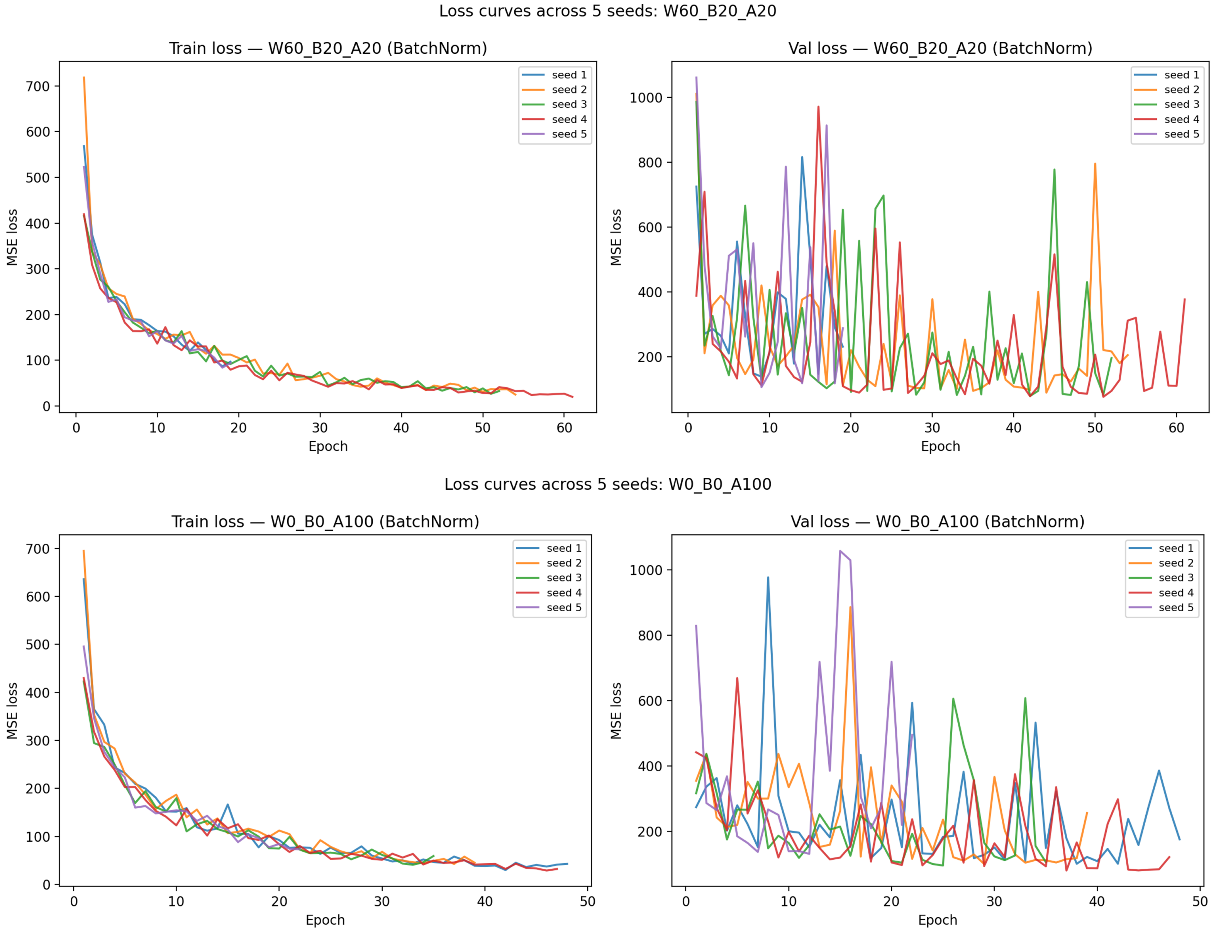


**Figure S1:** Per-epoch validation loss across five seeds under BatchNorm3D, for the 100% Asian and 60% W/ 20% B/20% A configurations at $n = 343$.

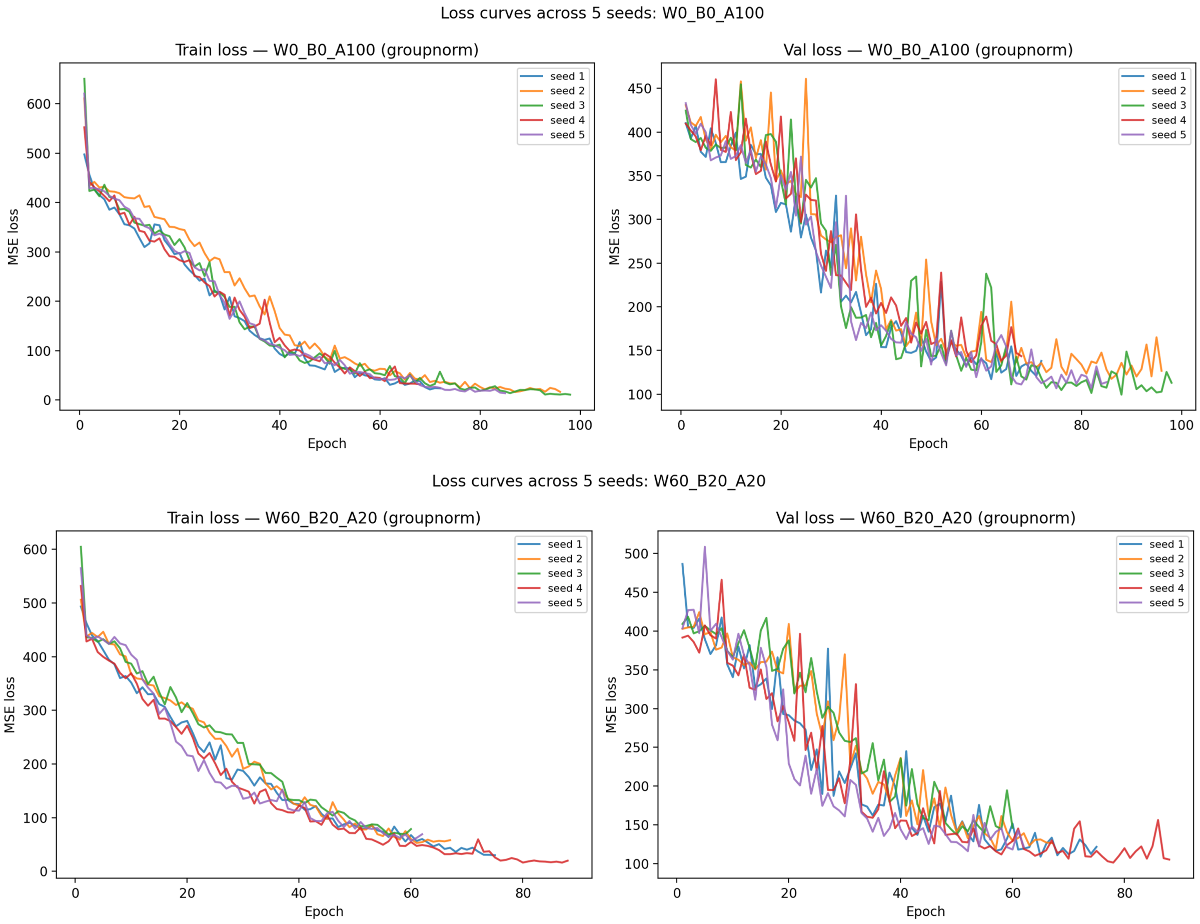


**Figure S2:** Per-epoch validation loss across five seeds under GroupNorm, for the 100% Asian and 60% W/ 20% B/20% A configurations at $n = 343$.

#### S2 Age-bias correction and BAG–age correlations

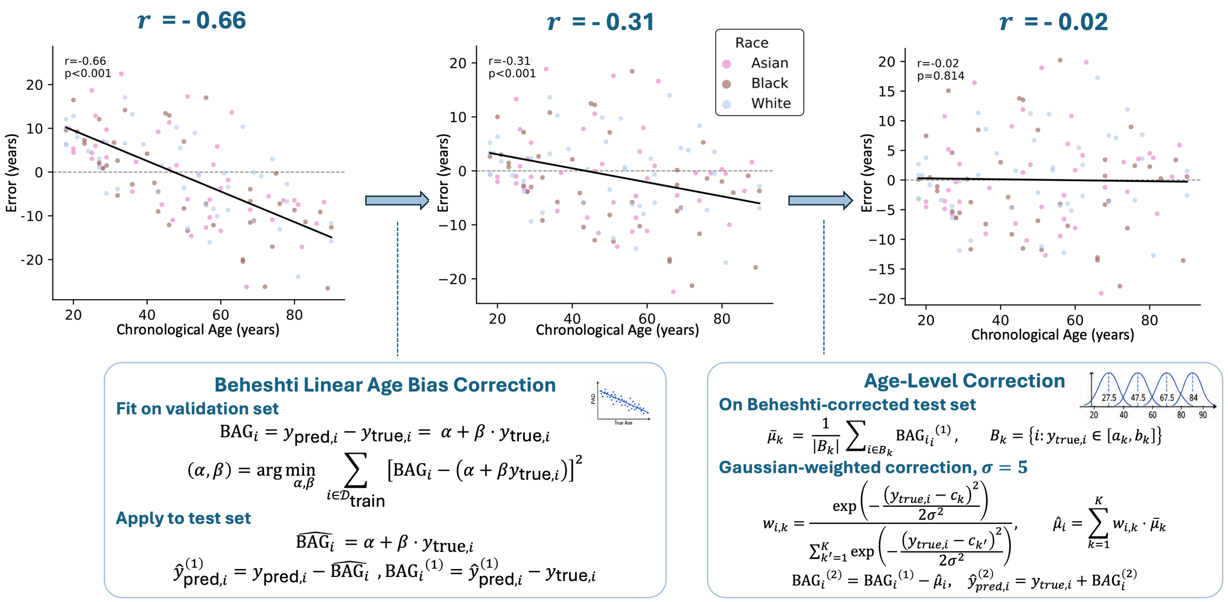


**Figure S3:** Example of the two-stage age-bias correction pipeline applied to the all-data model in the real domain. Brain age gap (BAG) is first corrected using a linear fit estimated on validation-set predictions and applied to test-set predictions, removing the bulk of the sample-level age bias. A residual soft-Gaussian age-bin correction is then applied directly to the Beheshti-corrected test predictions to drive any remaining age-structured BAG bias towards zero.

**Table S2:** BAG–age Pearson correlations at uncorrected, Beheshti-corrected, and final (soft-bin corrected) stages, pooled across race ($n = 132$), for all 13 training configurations. Raw $p$-values shown for the Beheshti stage; ten of thirteen real-domain and nine of thirteen synthetic-domain configurations retain a significant correlation at this stage, with all 26 cells reduced to non-significance ($p > 0.05$) after full correction.

| Configuration | Domain | $r_{\text{unc}}$ | $r_{\text{beh}}$ | $p_{\text{beh}}$ | Sig. | $r_{\text{final}}$ | $p_{\text{final}}$ |
|---|---|---|---|---|---|---|---|
| All-data | real | −0.543 | −0.205 | 0.0184 | * | −0.016 | 0.858 |
| 100% White (595W) | real | −0.664 | −0.313 | 0.0003 | *** | −0.021 | 0.814 |
| 100% Black (517B) | real | −0.678 | −0.229 | 0.0083 | ** | −0.015 | 0.867 |
| 100% Asian (343A) | real | −0.661 | −0.164 | 0.0610 | ns | −0.008 | 0.923 |
| 100% Black (343B) | real | −0.682 | −0.163 | 0.0620 | ns | −0.019 | 0.826 |
| 50% B/50% A | real | −0.565 | −0.172 | 0.0485 | * | −0.027 | 0.763 |
| 100% White (343W) | real | −0.635 | −0.222 | 0.0107 | * | −0.004 | 0.962 |
| 20% W/20% B/60% A | real | −0.600 | −0.255 | 0.0032 | ** | −0.018 | 0.838 |
| 20% W/60% B/20% A | real | −0.677 | −0.280 | 0.0011 | ** | −0.045 | 0.612 |
| 33%/33%/33% | real | −0.608 | −0.253 | 0.0034 | ** | −0.026 | 0.770 |
| 50% W/50% A | real | −0.621 | −0.226 | 0.0093 | ** | +0.004 | 0.960 |
| 50% W/50% B | real | −0.638 | −0.316 | 0.0002 | *** | −0.019 | 0.832 |
| 60% W/20% B/20% A | real | −0.591 | −0.166 | 0.0566 | ns | −0.022 | 0.803 |
| All-data | synth | −0.506 | −0.194 | 0.0261 | * | −0.020 | 0.820 |
| 100% White (595W) | synth | −0.565 | −0.241 | 0.0054 | ** | −0.039 | 0.658 |
| 100% Black (517B) | synth | −0.594 | −0.271 | 0.0016 | ** | −0.040 | 0.649 |
| 100% Asian (343A) | synth | −0.535 | −0.201 | 0.0210 | * | −0.035 | 0.690 |
| 100% Black (343B) | synth | −0.683 | −0.193 | 0.0266 | * | −0.036 | 0.682 |
| 50% B/50% A | synth | −0.585 | −0.226 | 0.0090 | ** | −0.016 | 0.853 |
| 100% White (343W) | synth | −0.556 | −0.098 | 0.2626 | ns | −0.016 | 0.860 |
| 20% W/20% B/60% A | synth | −0.526 | −0.212 | 0.0146 | * | −0.028 | 0.751 |
| 20% W/60% B/20% A | synth | −0.531 | −0.169 | 0.0529 | ns | −0.031 | 0.723 |
| 33%/33%/33% | synth | −0.567 | −0.242 | 0.0052 | ** | −0.037 | 0.672 |
| 50% W/50% A | synth | −0.513 | −0.187 | 0.0314 | * | −0.027 | 0.759 |
| 50% W/50% B | synth | −0.531 | −0.147 | 0.0934 | ns | −0.024 | 0.783 |
| 60% W/20% B/20% A | synth | −0.495 | −0.098 | 0.2622 | ns | −0.020 | 0.823 |

#### S3 Performance across all training configurations

**Table S3:** Real domain: per-race performance across all 13 training configurations. Uncorr $R^2$: coefficient of determination between raw predictions and chronological age per racial group. Linear MAE: Beheshti linear correction fitted on held-out validation set. Soft-bin MAE: additional soft-Gaussian age-bin correction on test set. RFG (fairness gap) = max(race MAE) − min(race MAE).

| | | | | **MAE (years)** | | **RFG** | |
|---|---|---|---|---|---|---|---|
| **Train** | **Test** | $N$ | **Uncorr** $R^2$ | **Linear** | **Soft-bin** | **Linear** | **Soft-bin** |
| **All (343A 517B 595W)** | Asian | 44 | 0.82 | $5.80 \pm 4.93$ | $5.70 \pm 4.82$ | 1.57 | 1.72 |
| | Black | 44 | 0.83 | $6.13 \pm 4.28$ | $5.91 \pm 4.20$ | | |
| | White | 44 | 0.89 | $4.56 \pm 3.18$ | $4.19 \pm 3.10$ | | |
| **100% Asian (343A)** | Asian | 44 | 0.60 | $8.15 \pm 5.99$ | $7.93 \pm 5.53$ | 2.03 | 2.42 |
| | Black | 44 | 0.61 | $8.23 \pm 6.60$ | $7.88 \pm 6.26$ | | |
| | White | 44 | 0.73 | $6.19 \pm 4.97$ | $5.51 \pm 4.17$ | | |
| **100% Black (517B)** | Asian | 44 | 0.63 | $7.18 \pm 6.04$ | $7.46 \pm 5.25$ | 1.45 | 1.88 |
| | Black | 44 | 0.73 | $6.61 \pm 5.46$ | $6.59 \pm 5.32$ | | |
| | White | 44 | 0.74 | $5.74 \pm 5.07$ | $5.59 \pm 4.23$ | | |
| **100% White (595W)** | Asian | 44 | 0.70 | $6.45 \pm 6.04$ | $6.70 \pm 5.24$ | 2.24 | 1.51 |
| | Black | 44 | 0.65 | $7.82 \pm 5.91$ | $7.07 \pm 5.36$ | | |
| | White | 44 | 0.81 | $5.58 \pm 4.22$ | $5.56 \pm 4.13$ | | |
| *Sweep configurations* ($n \sim 343$) | | | | | | | |
| **100% Black (343B)** | Asian | 44 | 0.52 | $9.10 \pm 6.27$ | $8.30 \pm 5.50$ | 3.03 | 2.30 |
| | Black | 44 | 0.65 | $7.12 \pm 6.65$ | $7.37 \pm 6.28$ | | |
| | White | 44 | 0.69 | $6.07 \pm 4.96$ | $6.00 \pm 4.94$ | | |
| **100% White (343W)** | Asian | 44 | 0.57 | $7.83 \pm 7.25$ | $8.10 \pm 6.29$ | 2.69 | 2.59 |
| | Black | 44 | 0.54 | $8.42 \pm 7.12$ | $7.85 \pm 6.30$ | | |
| | White | 44 | 0.77 | $5.73 \pm 4.36$ | $5.51 \pm 4.48$ | | |
| **60% W/20% B/20% A (206W 67B 68A)** | Asian | 44 | 0.61 | $7.96 \pm 6.40$ | $7.75 \pm 6.02$ | **3.00** | **2.88** |
| | Black | 44 | 0.58 | $8.80 \pm 6.60$ | $8.23 \pm 6.29$ | | |
| | White | 44 | 0.76 | $5.81 \pm 4.23$ | $5.35 \pm 3.77$ | | |
| **20% W/60% B/20% A (67W 206B 68A)** | Asian | 44 | 0.67 | $7.27 \pm 5.62$ | $7.43 \pm 5.35$ | **0.25** | **0.64** |
| | Black | 44 | 0.68 | $7.13 \pm 6.12$ | $6.96 \pm 5.74$ | | |
| | White | 44 | 0.68 | $7.02 \pm 5.28$ | $6.79 \pm 4.42$ | | |
| **20% W/20% B/60% A (67W 68B 206A)** | Asian | 44 | 0.61 | $8.34 \pm 6.50$ | $8.34 \pm 5.67$ | 1.55 | 2.00 |
| | Black | 44 | 0.62 | $7.65 \pm 7.59$ | $7.64 \pm 6.88$ | | |
| | White | 44 | 0.77 | $6.80 \pm 3.98$ | $6.35 \pm 4.12$ | | |
| **50% W/50% A (171W 172A)** | Asian | 44 | 0.61 | $7.93 \pm 6.74$ | $8.05 \pm 5.87$ | 2.55 | 2.73 |
| | Black | 44 | 0.67 | $7.07 \pm 6.57$ | $7.08 \pm 5.96$ | | |
| | White | 44 | 0.82 | $5.38 \pm 4.30$ | $5.32 \pm 4.38$ | | |
| **50% W/50% B (171W 172B)** | Asian | 44 | 0.49 | $9.58 \pm 7.48$ | $9.18 \pm 6.66$ | 2.61 | 2.55 |
| | Black | 44 | 0.62 | $7.61 \pm 7.26$ | $7.19 \pm 6.60$ | | |
| | White | 44 | 0.74 | $6.97 \pm 4.11$ | $6.63 \pm 3.96$ | | |
| **50% B/50% A (171B 172A)** | Asian | 44 | 0.61 | $8.74 \pm 6.57$ | $8.82 \pm 5.93$ | 2.51 | 2.70 |
| | Black | 44 | 0.64 | $7.57 \pm 7.28$ | $7.74 \pm 6.66$ | | |
| | White | 44 | 0.75 | $6.23 \pm 5.18$ | $6.13 \pm 4.88$ | | |
| **33% W/33% B/33% A (114W 114B 115A)** | Asian | 44 | 0.61 | $8.24 \pm 6.75$ | $8.21 \pm 6.07$ | 2.05 | 2.53 |
| | Black | 44 | 0.68 | $7.90 \pm 6.32$ | $7.43 \pm 6.00$ | | |
| | White | 44 | 0.77 | $6.19 \pm 4.70$ | $5.69 \pm 4.10$ | | |

In bold, the consistently worst and best RFG values after both linear and soft-bin correction.

**Table S4:** Synthetic domain: per-race performance across all 13 training configurations. Column definitions as in Table S3. Pareto-optimal configurations are marked ⋆.

| | | | | MAE (years) | | RFG | |
|---|---|---|---|---|---|---|---|
| **Train** | **Test** | $N$ | **Uncorr** $R^2$ | **Linear** | **Soft-bin** | **Linear** | **Soft-bin** |
| **All**⋆ | Asian | 44 | 0.84 | $5.09 \pm 4.96$ | $5.02 \pm 4.92$ | | |
| **(343A 517B 595W)** | Black | 44 | 0.86 | $5.81 \pm 3.75$ | $5.59 \pm 3.77$ | 0.72 | 0.80 |
| | White | 44 | 0.87 | $5.44 \pm 3.70$ | $4.78 \pm 3.47$ | | |
| **100% Asian** | Asian | 44 | 0.71 | $7.05 \pm 6.52$ | $6.15 \pm 6.16$ | | |
| **(343A)** | Black | 44 | 0.66 | $8.83 \pm 6.29$ | $6.86 \pm 5.76$ | 1.78 | 0.90 |
| | White | 44 | 0.77 | $7.19 \pm 4.95$ | $5.95 \pm 4.94$ | | |
| **100% Black** | Asian | 44 | 0.74 | $6.21 \pm 6.17$ | $6.16 \pm 5.83$ | | |
| **(517B)** | Black | 44 | 0.72 | $7.09 \pm 5.90$ | $6.78 \pm 5.54$ | 0.88 | 1.00 |
| | White | 44 | 0.79 | $6.61 \pm 4.41$ | $5.78 \pm 4.69$ | | |
| **100% White** | Asian | 44 | 0.72 | $6.32 \pm 6.47$ | $6.15 \pm 6.10$ | | |
| **(595W)** | Black | 44 | 0.68 | $7.65 \pm 6.46$ | $7.29 \pm 5.93$ | 1.33 | 1.41 |
| | White | 44 | 0.79 | $6.67 \pm 4.58$ | $5.88 \pm 4.84$ | | |
| *Sweep configurations* ($n \sim 343$) | | | | | | | |
| **100% Black** | Asian | 44 | 0.68 | $6.34 \pm 5.84$ | $6.03 \pm 5.58$ | | |
| **(343B)** | Black | 44 | 0.71 | $6.43 \pm 5.91$ | $6.46 \pm 5.86$ | 0.61 | 1.15 |
| | White | 44 | 0.75 | $5.82 \pm 4.74$ | $5.31 \pm 4.53$ | | |
| **100% White** | Asian | 44 | 0.69 | $7.38 \pm 6.21$ | $6.50 \pm 5.87$ | | |
| **(343W)** | Black | 44 | 0.65 | $8.26 \pm 5.68$ | $7.26 \pm 5.31$ | 2.79 | 1.71 |
| | White | 44 | 0.83 | $5.47 \pm 4.58$ | $5.56 \pm 4.97$ | | |
| **60% W/20% B/20% A** | Asian | 44 | 0.72 | $6.77 \pm 6.45$ | $6.40 \pm 6.30$ | | |
| **(206W 67B 68A)** | Black | 44 | 0.64 | $8.48 \pm 6.97$ | $8.33 \pm 6.74$ | **2.46** | **2.55** |
| | White | 44 | 0.81 | $6.02 \pm 5.20$ | $5.79 \pm 4.63$ | | |
| **20% W/60% B/20% A**⋆ | Asian | 44 | 0.76 | $6.05 \pm 6.07$ | $6.01 \pm 5.78$ | | |
| **(67W 206B 68A)** | Black | 44 | 0.76 | $6.14 \pm 6.42$ | $6.52 \pm 6.02$ | **0.21** | **0.59** |
| | White | 44 | 0.80 | $6.26 \pm 4.82$ | $5.93 \pm 4.32$ | | |
| **20% W/20% B/60% A** | Asian | 44 | 0.73 | $6.84 \pm 6.33$ | $6.28 \pm 6.10$ | | |
| **(67W 68B 206A)** | Black | 44 | 0.67 | $8.37 \pm 6.73$ | $7.70 \pm 6.71$ | 2.04 | 1.87 |
| | White | 44 | 0.82 | $6.33 \pm 4.32$ | $5.83 \pm 4.51$ | | |
| **50% W/50% A** | Asian | 44 | 0.72 | $7.17 \pm 6.33$ | $6.75 \pm 6.09$ | | |
| **(171W 172A)** | Black | 44 | 0.61 | $9.23 \pm 6.84$ | $8.30 \pm 6.40$ | 2.82 | 1.73 |
| | White | 44 | 0.81 | $6.41 \pm 5.19$ | $6.57 \pm 5.42$ | | |
| **50% W/50% B** | Asian | 44 | 0.69 | $6.57 \pm 6.93$ | $6.63 \pm 6.29$ | | |
| **(171W 172B)** | Black | 44 | 0.71 | $7.10 \pm 6.30$ | $6.64 \pm 6.30$ | 1.21 | 1.04 |
| | White | 44 | 0.82 | $5.89 \pm 4.04$ | $5.60 \pm 3.87$ | | |
| **50% B/50% A** | Asian | 44 | 0.75 | $6.26 \pm 5.91$ | $6.22 \pm 5.59$ | | |
| **(171B 172A)** | Black | 44 | 0.75 | $6.22 \pm 6.40$ | $6.30 \pm 5.93$ | 0.27 | 0.82 |
| | White | 44 | 0.81 | $5.99 \pm 4.26$ | $5.48 \pm 3.68$ | | |
| **33% W/33% B/33% A** | Asian | 44 | 0.74 | $6.46 \pm 6.03$ | $6.46 \pm 5.64$ | | |
| **(114W 114B 115A)** | Black | 44 | 0.69 | $7.60 \pm 6.62$ | $7.39 \pm 6.41$ | 1.49 | 1.72 |
| | White | 44 | 0.81 | $6.11 \pm 4.58$ | $5.67 \pm 4.39$ | | |

⋆ Pareto-optimal: not dominated on both overall MAE and RFG simultaneously by any configuration in either domain. All-data synthetic maximizes accuracy; 20% W/60% B/20% A synthetic maximizes fairness.
In bold, the consistently worst and best RFG values after both linear and soft-bin correction.

### S4 External benchmark models

**Table S5:** Per-race performance for BrainAgeNeXt and SynthBA evaluated on the fixed balanced internal test set ($N = 44$ per race) in both real and synthetic domains. MAE is reported uncorrected, after linear (Beheshti) correction, and after the additional soft-Gaussian bin correction; RFG is the corresponding max−min per-race MAE at each stage. Bold RFG values indicate the lowest fairness gap achieved for each model across imaging domains.

| | | | | MAE (years) | | | RFG | | |
|---|---|---|---|---|---|---|---|---|---|
| **Model** | **Test** | $N$ | **Uncorr** $R^2$ | **Uncorr** | **Linear** | **Soft-bin** | **Uncorr** | **Linear** | **Soft-bin** |
| **BrainAgeNeXt (Real)** | Asian | 44 | 0.912 | $4.35 \pm 3.97$ | $4.15 \pm 4.06$ | $4.08 \pm 4.09$ | 1.79 | 1.30 | 0.82 |
| | Black | 44 | 0.880 | $6.15 \pm 4.10$ | $5.45 \pm 3.89$ | $4.90 \pm 3.79$ | | | |
| | White | 44 | 0.925 | $4.72 \pm 3.37$ | $4.66 \pm 2.92$ | $4.25 \pm 3.21$ | | | |
| **BrainAgeNeXt (Synthetic)** | Asian | 44 | 0.785 | $7.44 \pm 5.41$ | $5.14 \pm 5.00$ | $4.94 \pm 5.08$ | **1.42** | **0.55** | **0.49** |
| | Black | 44 | 0.763 | $8.29 \pm 6.22$ | $5.66 \pm 4.54$ | $5.17 \pm 4.23$ | | | |
| | White | 44 | 0.819 | $6.87 \pm 5.78$ | $5.11 \pm 4.01$ | $4.68 \pm 3.77$ | | | |
| **SynthBA (Real)** | Asian | 44 | 0.129 | $15.17 \pm 10.59$ | $8.93 \pm 9.33$ | $8.56 \pm 8.98$ | 4.13 | 1.33 | 0.96 |
| | Black | 44 | 0.197 | $15.96 \pm 10.46$ | $8.98 \pm 8.21$ | $8.53 \pm 7.06$ | | | |
| | White | 44 | 0.487 | $11.83 \pm 9.47$ | $7.65 \pm 6.91$ | $7.60 \pm 6.51$ | | | |
| **SynthBA (Synthetic)** | Asian | 44 | 0.213 | $14.73 \pm 9.61$ | $8.51 \pm 8.87$ | $7.76 \pm 8.88$ | **2.87** | **0.64** | **0.81** |
| | Black | 44 | 0.341 | $14.12 \pm 9.98$ | $8.25 \pm 7.69$ | $8.35 \pm 6.63$ | | | |
| | White | 44 | 0.478 | $11.87 \pm 9.62$ | $7.87 \pm 6.98$ | $7.54 \pm 6.14$ | | | |

### S5 Statistical significance of racial and sex fairness gaps

#### Within-model tests

Within each of the 26 model × domain cells, race (three levels: Asian, Black, White; $n = 44$ per group), sex (Female vs Male), and sex × race (six levels, $n = 16$–28 per group) subgroup effects on BAG were each assessed via the Kruskal–Wallis test, reported as the epsilon-squared effect size ($\varepsilon^2$), at the Beheshti-corrected stage (Table S6). $p$-values were FDR-corrected (Benjamini–Hochberg) separately within each of the three 26-cell test families. No cell survives FDR correction for any of the three factors. The closest approach to significance at the raw level was a sex effect in the 20% W/60% B/20% A configuration, real domain ($p = 0.0263$, $\varepsilon^2 = 0.030$), which did not survive FDR correction ($p_{\mathrm{FDR}} = 0.310$); the closest race effect was 100% Black (343B), real domain ($p = 0.0601$, $\varepsilon^2 = 0.028$, $p_{\mathrm{FDR}} = 0.781$); the closest sex × race effect was the same configuration ($p = 0.0856$, $\varepsilon^2 = 0.037$, $p_{\mathrm{FDR}} = 0.433$). This uniformly null result reflects a power limitation rather than an absence of effect: with only 44 subjects per racial group and 16–28 per sex × race cell, within-model tests are underpowered to detect the effect sizes observed at the model level (main text, Statistical Analysis).

**Table S6:** Within-model race, sex, and sex × race subgroups effects on BAG (Kruskal–Wallis, $\varepsilon^2$) at the Beheshti-corrected stage, all 26 model × domain cells. $p_{\mathrm{FDR}}$: Benjamini–Hochberg correction applied separately within each of the three 26-cell test families. No cell is significant after correction for any factor.

| Configuration | Domain | $\varepsilon^2_{\mathrm{race}}$ | $p_{\mathrm{FDR,race}}$ | $\varepsilon^2_{\mathrm{sex}}$ | $p_{\mathrm{FDR,sex}}$ | $\varepsilon^2_{\mathrm{sex}\times\mathrm{race}}$ | $p_{\mathrm{FDR,sex}\times\mathrm{race}}$ |
|---|---|---|---|---|---|---|---|
| All-data | real | 0.0000 | 0.903 | 0.0110 | 0.310 | 0.0158 | 0.720 |
| 100% White (595W) | real | 0.0000 | 0.903 | 0.0054 | 0.357 | 0.0000 | 0.886 |
| 100% Black (517B) | real | 0.0168 | 0.809 | 0.0193 | 0.310 | 0.0336 | 0.433 |
| 100% Asian (343A) | real | 0.0000 | 0.903 | 0.0000 | 0.628 | 0.0000 | 0.886 |
| 100% Black (343B) | real | 0.0281 | 0.781 | 0.0173 | 0.310 | 0.0370 | 0.433 |
| 50% B/50% A | real | 0.0000 | 0.903 | 0.0000 | 0.958 | 0.0000 | 0.913 |
| 100% White (343W) | real | 0.0000 | 0.903 | 0.0007 | 0.428 | 0.0000 | 0.886 |
| 20% W/20% B/60% A | real | 0.0000 | 0.903 | 0.0002 | 0.428 | 0.0000 | 0.886 |
| 20% W/60% B/20% A | real | 0.0032 | 0.903 | 0.0303 | 0.310 | 0.0345 | 0.433 |
| 33%/33%/33% | real | 0.0000 | 0.903 | 0.0119 | 0.310 | 0.0037 | 0.886 |
| 50% W/50% A | real | 0.0000 | 0.903 | 0.0000 | 0.799 | 0.0000 | 0.886 |
| 50% W/50% B | real | 0.0000 | 0.903 | 0.0000 | 0.428 | 0.0000 | 0.886 |
| 60% W/20% B/20% A | real | 0.0000 | 0.903 | 0.0056 | 0.357 | 0.0000 | 0.886 |
| All-data | synth | 0.0000 | 0.950 | 0.0069 | 0.434 | 0.0132 | 0.599 |
| 100% White (595W) | synth | 0.0168 | 0.444 | 0.0025 | 0.434 | 0.0088 | 0.599 |
| 100% Black (517B) | synth | 0.0000 | 0.795 | 0.0005 | 0.434 | 0.0000 | 0.599 |
| 100% Asian (343A) | synth | 0.0173 | 0.444 | 0.0008 | 0.434 | 0.0085 | 0.599 |
| 100% Black (343B) | synth | 0.0026 | 0.795 | 0.0128 | 0.434 | 0.0120 | 0.599 |
| 50% B/50% A | synth | 0.0000 | 0.982 | 0.0000 | 0.434 | 0.0000 | 0.599 |
| 100% White (343W) | synth | 0.0168 | 0.444 | 0.0000 | 0.784 | 0.0118 | 0.599 |
| 20% W/20% B/60% A | synth | 0.0000 | 0.950 | 0.0000 | 0.529 | 0.0000 | 0.689 |
| 20% W/60% B/20% A | synth | 0.0000 | 0.950 | 0.0068 | 0.434 | 0.0000 | 0.599 |
| 33%/33%/33% | synth | 0.0000 | 0.950 | 0.0039 | 0.434 | 0.0069 | 0.599 |
| 50% W/50% A | synth | 0.0154 | 0.444 | 0.0000 | 0.556 | 0.0021 | 0.599 |
| 50% W/50% B | synth | 0.0000 | 0.982 | 0.0008 | 0.434 | 0.0000 | 0.620 |
| 60% W/20% B/20% A | synth | 0.0000 | 0.950 | 0.0039 | 0.434 | 0.0000 | 0.599 |

#### Across-model tests: race gap versus sex gap

Treating each of the 13 training configurations as a single observation, Wilcoxon signed-rank tests (paired on configuration identity) compared the racial fairness gap against the sex fairness gap at each correction stage and imaging domain. At the Beheshti-corrected stage, race disparities exceeded sex disparities in 12 of 13 real-domain configurations (paired Wilcoxon W=4, p=0.0017), with mean race and sex gaps of 2.12 and 1.07 years, respectively. In the synthetic domain, the two disparities converged: the race gap exceeded the sex gap in 6 of 13 configurations, and the paired difference was not statistically significant (W=38, p=0.636); mean race and sex gaps were 1.43 and 1.27 years, respectively. Switching to synthetic imaging significantly reduced the race gap, whereas the sex gap did not change significantly (1.07 to 1.27 years; W=19, p=0.068). The non-significance at the synthetic Beheshti stage reflects a genuine convergence of race and sex disparities in this domain rather than a statistical artifact of reduced power, since the same 13-configuration design achieves $r = 1.0$ in the real domain.

#### Cross-domain paired tests

Paired Wilcoxon signed-rank tests compared real- and synthetic-domain BAG for the same subjects within each model, separately by race, yielding 39 model × race comparisons (13 models × 3 racial groups), each based on 44 paired subjects, with FDR correction applied within the test family at the Beheshti-corrected stage. 19 of 39 comparisons were significant after FDR correction ($p_{\text{FDR}} < 0.05$). The domain shift was racially asymmetric: Asian subjects showed the largest median shift in BAG between domains (synthetic minus real, median across the 13 configurations: $-1.73$ years, significant in 8/13 configurations), followed by Black subjects ($-0.99$ years, 7/13 significant), while White subjects showed a smaller shift ($-0.33$ years, 4/13 significant), indicating that the synthetic domain's effect on BAG is concentrated in the groups most disadvantaged in the real domain.

This asymmetric compression translates directly into a lower model-level racial fairness gap in the synthetic domain. Treating each configuration's racial RFG as a paired model-level observation, racial RFG was significantly lower in the synthetic domain after Beheshti correction (mean 1.43 vs 2.12 years; median 1.33 vs 2.24 years; W=11, p=0.0134) and after full correction (mean 1.33 vs 2.19 years; median 1.15 vs 2.42 years; W=0, p=0.000244). This is consistent with the per-race result above, in which Asian and Black subjects (the two groups with the largest real-domain BAG) show the largest synthetic-domain reduction, and confirms that the domain effect on racial disparity is not an artifact of a single correction stage.

#### S6 Race × sex subgroup performance

##### Methods

Race × sex subgroup performance was assessed for all 13 training configurations in both imaging domains by stratifying the balanced test set ($n = 44$ per race) into six race × sex cells. Cell sizes reflect the underlying sex distribution within each racial group in the test set (Asian: 22F/22M; Black: 27F/17M; White: 28F/16M); because the smallest cells contain 16–17 subjects, per-cell estimates are correspondingly less stable than the race-level estimates reported in the main text. $R^2$ was computed on uncorrected predictions per cell; MAE is reported after linear (Beheshti) correction and after full soft-bin correction. The sex × race fairness gap (RFG) is the maximum minus minimum subgroup MAE across all six cells within each model, reported at both correction stages.

##### Results

Tables S7 and S8 report per-cell performance for the real and synthetic domains, respectively, across all 13 training configurations.

At the Beheshti-corrected stage, race disparities exceeded sex disparities in 12 of 13 real-domain configurations (W=4, p=0.0017), whereas in the synthetic domain the race and sex gaps did not differ significantly (W=38, p=0.636). Switching to synthetic imaging reduced race disparities more strongly than sex disparities: the mean Beheshti-corrected race gap decreased from 2.12 to 1.43 years (32% reduction), whereas the corresponding sex gap remained similar (1.07 to 1.28 years).

Joint sex-and-race analysis revealed larger disparities than race-only analyses, and the sex × race RFG exceeded the corresponding race-only RFG in every configuration, reflecting the more extreme performance of the single worst subgroup relative to the racial group mean. Across configurations, the Black–White MAE difference was larger in males than in females (1.73 vs 1.50 years), and the Asian–White difference was more than twice as large in males (2.75 vs 1.18 years).

The identity of the worst-predicted subgroup differed between domains. In the real domain, Asian Male and Black Female subjects accounted for the majority of worst-performing cells: Asian Male was the worst subgroup in 7 of 13 configurations, reaching MAE = 10.56 years in the fixed-size 100% Black ($n = 343$) model, while Black Female was worst in 3 of 13. In the synthetic domain, Black Female consolidated as the worst-predicted subgroup in every configuration without exception (MAE range 6.26–9.88 years). The combined sex × race fairness gap was nonetheless lower in the synthetic domain in 9 of 13 configurations after Beheshti correction.

**Table S7:** Race × sex subgroup performance, real MRI domain. $R^2$ before correction; MAE after linear and after final soft-bin correction. $\mathrm{RFG}_{\mathrm{linear}}$ / $\mathrm{RFG}_{\mathrm{softbin}}$: max−min subgroup MAE across the six race × sex cells.

| Train | Test Group | N | $R^2_{\mathrm{uncorr}}$ | $\mathrm{MAE}_{\mathrm{linear}}$ (± std) | $\mathrm{MAE}_{\mathrm{softbin}}$ (± std) | $\mathrm{RFG}_{\mathrm{lin}}$ / $\mathrm{RFG}_{\mathrm{soft}}$ |
|---|---|---|---|---|---|---|
| **All-data** | Asian Female | 22 | 0.85 | 5.33 ± 4.25 | 4.95 ± 3.70 | |
| | Asian Male | 22 | 0.74 | 6.27 ± 5.69 | 6.45 ± 5.81 | |
| | Black Female | 27 | 0.83 | 6.01 ± 4.36 | 5.87 ± 4.30 | |
| | Black Male | 17 | 0.84 | 6.32 ± 4.40 | 5.96 ± 4.29 | |
| | White Female | 28 | 0.88 | 4.95 ± 3.39 | 4.26 ± 3.42 | |
| | White Male | 16 | 0.92 | 3.86 ± 2.87 | 4.05 ± 2.67 | 2.45 / 2.40 |
| **100% White (595W)** | Asian Female | 22 | 0.73 | 5.67 ± 6.28 | 5.60 ± 5.10 | |
| | Asian Male | 22 | 0.63 | 7.23 ± 5.96 | 7.80 ± 5.38 | |
| | Black Female | 27 | 0.58 | 8.32 ± 6.37 | 7.84 ± 5.56 | |
| | Black Male | 17 | 0.73 | 7.02 ± 5.39 | 5.84 ± 5.11 | |
| | White Female | 28 | 0.81 | 5.92 ± 4.11 | 5.75 ± 4.19 | |
| | White Male | 16 | 0.81 | 4.99 ± 4.62 | 5.23 ± 4.26 | 3.33 / 2.61 |
| **100% White (343W)** | Asian Female | 22 | 0.60 | 8.22 ± 7.37 | 8.09 ± 6.46 | |
| | Asian Male | 22 | 0.51 | 7.96 ± 6.84 | 8.54 ± 6.21 | |
| | Black Female | 27 | 0.48 | 8.40 ± 7.98 | 8.27 ± 7.08 | |
| | Black Male | 17 | 0.62 | 8.72 ± 6.48 | 8.18 ± 5.86 | |
| | White Female | 28 | 0.80 | 5.67 ± 4.74 | 5.68 ± 5.28 | |
| | White Male | 16 | 0.78 | 5.12 ± 4.23 | 5.28 ± 3.68 | 3.61 / 3.25 |
| **100% Black (517B)** | Asian Female | 22 | 0.66 | 7.19 ± 6.13 | 7.49 ± 4.39 | |
| | Asian Male | 22 | 0.54 | 7.18 ± 6.24 | 7.43 ± 6.19 | |
| | Black Female | 27 | 0.67 | 7.45 ± 5.97 | 7.69 ± 5.46 | |
| | Black Male | 17 | 0.79 | 5.26 ± 4.58 | 4.83 ± 4.92 | |
| | White Female | 28 | 0.70 | 6.13 ± 5.73 | 5.46 ± 4.93 | |
| | White Male | 16 | 0.82 | 5.06 ± 3.96 | 5.81 ± 2.96 | 2.40 / 2.86 |
| **100% Black (343B)** | Asian Female | 22 | 0.61 | 8.27 ± 6.11 | 7.75 ± 5.00 | |
| | Asian Male | 22 | 0.27 | 10.56 ± 6.97 | 8.69 ± 6.15 | |
| | Black Female | 27 | 0.58 | 8.01 ± 7.55 | 8.38 ± 6.99 | |
| | Black Male | 17 | 0.73 | 5.92 ± 5.45 | 5.53 ± 4.92 | |
| | White Female | 28 | 0.65 | 6.38 ± 5.34 | 6.09 ± 5.18 | |
| | White Male | 16 | 0.77 | 5.86 ± 4.30 | 5.63 ± 4.71 | 4.70 / 3.16 |
| **100% Asian (343A)** | Asian Female | 22 | 0.66 | 7.77 ± 5.77 | 7.33 ± 4.66 | |
| | Asian Male | 22 | 0.46 | 8.33 ± 6.60 | 8.76 ± 6.14 | |
| | Black Female | 27 | 0.63 | 7.96 ± 6.60 | 7.84 ± 6.35 | |
| | Black Male | 17 | 0.59 | 8.22 ± 7.28 | 8.12 ± 6.42 | |
| | White Female | 28 | 0.73 | 6.64 ± 4.58 | 5.71 ± 4.28 | |
| | White Male | 16 | 0.76 | 4.47 ± 4.62 | 4.31 ± 3.46 | 3.85 / 4.44 |
| **20% W/20% B/60% A** | Asian Female | 22 | 0.69 | 7.72 ± 6.19 | 7.66 ± 5.30 | |
| | Asian Male | 22 | 0.40 | 9.20 ± 7.59 | 9.03 ± 6.34 | |
| | Black Female | 27 | 0.57 | 7.88 ± 8.24 | 8.04 ± 6.97 | |
| | Black Male | 17 | 0.67 | 7.88 ± 6.98 | 7.36 ± 7.19 | |
| | White Female | 28 | 0.78 | 6.86 ± 4.17 | 6.29 ± 4.31 | |
| | White Male | 16 | 0.76 | 6.56 ± 4.18 | 6.30 ± 3.62 | 2.64 / 2.74 |
| **50% B/50% A** | Asian Female | 22 | 0.71 | 8.19 ± 5.42 | 7.86 ± 5.31 | |
| | Asian Male | 22 | 0.43 | 9.31 ± 7.81 | 9.67 ± 6.84 | |
| | Black Female | 27 | 0.66 | 7.06 ± 7.13 | 7.34 ± 6.40 | |
| | Black Male | 17 | 0.64 | 8.60 ± 7.46 | 8.64 ± 7.22 | |
| | White Female | 28 | 0.77 | 6.00 ± 5.02 | 5.74 ± 4.63 | |
| | White Male | 16 | 0.76 | 6.19 ± 5.60 | 6.64 ± 5.29 | 3.31 / 3.92 |
| **50% W/50% A** | Asian Female | 22 | 0.68 | 7.34 ± 6.02 | 7.32 ± 5.08 | |
| | Asian Male | 22 | 0.50 | 8.19 ± 7.34 | 8.34 ± 6.58 | |
| | Black Female | 27 | 0.70 | 6.09 ± 6.71 | 6.27 ± 5.91 | |
| | Black Male | 17 | 0.66 | 8.16 ± 6.22 | 7.82 ± 6.12 | |
| | White Female | 28 | 0.83 | 5.24 ± 4.17 | 4.99 ± 4.32 | |
| | White Male | 16 | 0.80 | 5.98 ± 4.54 | 6.18 ± 4.41 | 2.94 / 3.35 |
| **33% W/33% B/33% A** | Asian Female | 22 | 0.69 | 7.56 ± 6.42 | 7.03 ± 4.50 | |
| | Asian Male | 22 | 0.40 | 9.49 ± 7.50 | 9.51 ± 7.40 | |
| | Black Female | 27 | 0.69 | 8.00 ± 5.74 | 7.12 ± 6.28 | |
| | Black Male | 17 | 0.63 | 8.89 ± 7.36 | 7.81 ± 6.24 | |

*continued on next page*

*Table S7 (continued)*

| Train | Test Group | N | $R^2_{\text{uncorr}}$ | $\text{MAE}_{\text{linear}}$ (± std) | $\text{MAE}_{\text{softbin}}$ (± std) | $\text{RFG}_{\text{lin}}$ / $\text{RFG}_{\text{soft}}$ |
|---|---|---|---|---|---|---|
| | White Female | 28 | 0.74 | 7.03 ± 4.95 | 5.07 ± 3.88 | |
| | White Male | 16 | 0.74 | 6.02 ± 6.01 | 7.04 ± 4.87 | 3.47 / 4.43 |
| **20% W/60% B/20% A** | Asian Female | 22 | 0.72 | 6.62 ± 5.49 | 6.33 ± 4.10 | |
| | Asian Male | 22 | 0.56 | 7.81 ± 6.16 | 8.44 ± 6.51 | |
| | Black Female | 27 | 0.63 | 7.56 ± 6.52 | 7.08 ± 6.42 | |
| | Black Male | 17 | 0.68 | 7.12 ± 6.62 | 6.90 ± 5.10 | |
| | White Female | 28 | 0.62 | 7.90 ± 5.85 | 6.83 ± 4.20 | |
| | White Male | 16 | 0.68 | 6.22 ± 6.43 | 6.68 ± 5.81 | 1.69 / 2.11 |
| **60% W/20% B/20% A** | Asian Female | 22 | 0.70 | 7.56 ± 5.70 | 7.33 ± 5.48 | |
| | Asian Male | 22 | 0.46 | 8.85 ± 7.49 | 8.91 ± 6.99 | |
| | Black Female | 27 | 0.55 | 9.04 ± 7.02 | 9.19 ± 6.13 | |
| | Black Male | 17 | 0.61 | 8.61 ± 7.69 | 8.34 ± 7.79 | |
| | White Female | 28 | 0.81 | 5.23 ± 3.92 | 5.25 ± 3.92 | |
| | White Male | 16 | 0.75 | 6.77 ± 4.42 | 6.56 ± 4.08 | 3.81 / 3.94 |
| **50% W/50% B** | Asian Female | 22 | 0.58 | 8.67 ± 7.55 | 8.47 ± 6.58 | |
| | Asian Male | 22 | 0.28 | 10.35 ± 8.05 | 9.88 ± 7.09 | |
| | Black Female | 27 | 0.53 | 8.51 ± 7.60 | 7.65 ± 6.97 | |
| | Black Male | 17 | 0.70 | 6.79 ± 6.68 | 6.10 ± 6.25 | |
| | White Female | 28 | 0.75 | 6.85 ± 3.66 | 6.28 ± 3.67 | |
| | White Male | 16 | 0.67 | 7.89 ± 4.91 | 7.54 ± 4.63 | 3.56 / 3.79 |

**Table S8:** Race × sex subgroup performance, synthetic T1 domain. Notation as in Table S7.

| Train | Test Group | N | $R^2_{\text{uncorr}}$ | $\text{MAE}_{\text{linear}}$ (± std) | $\text{MAE}_{\text{softbin}}$ (± std) | $\text{RFG}_{\text{lin}}$ / $\text{RFG}_{\text{soft}}$ |
|---|---|---|---|---|---|---|
| **All-data** | Asian Female | 22 | 0.87 | 5.32 ± 3.71 | 4.68 ± 3.17 | |
| | Asian Male | 22 | 0.79 | 4.86 ± 6.15 | 5.36 ± 6.35 | |
| | Black Female | 27 | 0.83 | 6.26 ± 3.96 | 6.18 ± 3.93 | |
| | Black Male | 17 | 0.90 | 5.09 ± 3.50 | 4.65 ± 3.54 | |
| | White Female | 28 | 0.87 | 5.68 ± 3.00 | 4.77 ± 2.98 | |
| | White Male | 16 | 0.85 | 5.03 ± 4.87 | 4.80 ± 4.40 | 1.40 / 1.53 |
| **100% White (595W)** | Asian Female | 22 | 0.72 | 6.83 ± 6.36 | 6.26 ± 5.22 | |
| | Asian Male | 22 | 0.70 | 5.82 ± 6.84 | 6.05 ± 7.12 | |
| | Black Female | 27 | 0.57 | 8.75 ± 7.08 | 8.52 ± 6.26 | |
| | Black Male | 17 | 0.81 | 5.91 ± 5.28 | 5.34 ± 5.15 | |
| | White Female | 28 | 0.78 | 6.86 ± 4.57 | 6.09 ± 4.56 | |
| | White Male | 16 | 0.80 | 6.33 ± 4.87 | 5.51 ± 5.57 | 2.93 / 3.18 |
| **100% White (343W)** | Asian Female | 22 | 0.66 | 8.12 ± 6.94 | 6.87 ± 6.28 | |
| | Asian Male | 22 | 0.75 | 6.12 ± 5.50 | 6.23 ± 5.86 | |
| | Black Female | 27 | 0.62 | 8.29 ± 5.36 | 7.50 ± 5.47 | |
| | Black Male | 17 | 0.72 | 7.82 ± 5.91 | 7.02 ± 5.59 | |
| | White Female | 28 | 0.87 | 4.93 ± 4.00 | 5.07 ± 4.33 | |
| | White Male | 16 | 0.78 | 5.92 ± 5.91 | 6.49 ± 6.01 | 3.36 / 2.43 |
| **100% Black (517B)** | Asian Female | 22 | 0.74 | 6.46 ± 5.95 | 5.95 ± 5.13 | |
| | Asian Male | 22 | 0.70 | 5.96 ± 6.65 | 6.36 ± 6.69 | |
| | Black Female | 27 | 0.65 | 7.75 ± 6.21 | 7.40 ± 5.69 | |
| | Black Male | 17 | 0.80 | 6.06 ± 5.60 | 5.78 ± 5.48 | |
| | White Female | 28 | 0.78 | 6.90 ± 4.38 | 5.88 ± 4.57 | |
| | White Male | 16 | 0.81 | 6.09 ± 4.70 | 5.60 ± 5.18 | 1.78 / 1.80 |
| **100% Black (343B)** | Asian Female | 22 | 0.73 | 5.83 ± 4.94 | 5.21 ± 4.37 | |
| | Asian Male | 22 | 0.59 | 6.57 ± 6.75 | 6.51 ± 6.79 | |
| | Black Female | 27 | 0.62 | 7.48 ± 6.53 | 7.56 ± 6.23 | |
| | Black Male | 17 | 0.81 | 4.61 ± 4.92 | 4.65 ± 5.24 | |
| | White Female | 28 | 0.72 | 6.21 ± 4.93 | 5.68 ± 4.54 | |
| | White Male | 16 | 0.80 | 4.98 ± 4.55 | 4.71 ± 4.59 | 2.87 / 2.91 |
| **100% Asian (343A)** | Asian Female | 22 | 0.70 | 7.50 ± 6.81 | 6.36 ± 5.06 | |
| | Asian Male | 22 | 0.72 | 6.28 ± 6.54 | 6.03 ± 7.37 | |
| | Black Female | 27 | 0.57 | 9.55 ± 6.58 | 7.88 ± 6.00 | |
| | Black Male | 17 | 0.77 | 7.55 ± 6.09 | 5.58 ± 5.74 | |
| | White Female | 28 | 0.77 | 7.46 ± 4.76 | 6.22 ± 4.61 | |
| | White Male | 16 | 0.79 | 6.71 ± 5.32 | 5.82 ± 5.89 | 3.27 / 2.29 |
| **20% W/20% B/60% A** | Asian Female | 22 | 0.72 | 7.71 ± 6.29 | 6.56 ± 5.05 | |
| | Asian Male | 22 | 0.67 | 6.56 ± 6.81 | 6.07 ± 7.39 | |
| | Black Female | 27 | 0.58 | 9.24 ± 6.93 | 8.46 ± 7.11 | |
| | Black Male | 17 | 0.74 | 7.68 ± 7.02 | 6.73 ± 6.59 | |
| | White Female | 28 | 0.80 | 6.80 ± 4.75 | 5.97 ± 4.54 | |
| | White Male | 16 | 0.83 | 5.97 ± 4.19 | 5.83 ± 4.84 | 3.27 / 2.63 |
| **50% B/50% A** | Asian Female | 22 | 0.77 | 6.36 ± 5.44 | 6.11 ± 5.25 | |
| | Asian Male | 22 | 0.67 | 6.31 ± 6.74 | 6.51 ± 6.22 | |
| | Black Female | 27 | 0.65 | 7.18 ± 7.45 | 7.23 ± 6.69 | |
| | Black Male | 17 | 0.85 | 4.64 ± 4.58 | 4.74 ± 4.62 | |
| | White Female | 28 | 0.80 | 6.31 ± 4.46 | 5.80 ± 3.82 | |
| | White Male | 16 | 0.82 | 5.38 ± 4.13 | 4.84 ± 3.69 | 2.54 / 2.50 |
| **50% W/50% A** | Asian Female | 22 | 0.70 | 7.49 ± 6.60 | 6.76 ± 5.50 | |
| | Asian Male | 22 | 0.70 | 6.81 ± 6.39 | 6.87 ± 6.87 | |
| | Black Female | 27 | 0.52 | 9.88 ± 6.55 | 9.00 ± 6.03 | |
| | Black Male | 17 | 0.71 | 7.79 ± 7.59 | 6.90 ± 7.11 | |
| | White Female | 28 | 0.82 | 6.20 ± 5.03 | 6.30 ± 5.09 | |
| | White Male | 16 | 0.76 | 7.07 ± 5.53 | 7.22 ± 5.96 | 3.68 / 2.70 |
| **33% W/33% B/33% A** | Asian Female | 22 | 0.75 | 6.85 ± 5.74 | 6.57 ± 4.87 | |
| | Asian Male | 22 | 0.68 | 6.33 ± 6.52 | 6.53 ± 6.52 | |
| | Black Female | 27 | 0.60 | 8.34 ± 7.22 | 8.24 ± 6.78 | |
| | Black Male | 17 | 0.79 | 6.33 ± 5.65 | 5.88 ± 5.79 | |
| | White Female | 28 | 0.81 | 6.37 ± 4.15 | 5.69 ± 4.14 | |
| | White Male | 16 | 0.78 | 5.87 ± 5.50 | 5.79 ± 5.17 | 2.47 / 2.54 |

*continued on next page*

*Table S8 (continued)*

| Train | Test Group | N | $R^2_{\text{uncorr}}$ | $\text{MAE}_{\text{linear}}$ ($\pm$ std) | $\text{MAE}_{\text{softbin}}$ ($\pm$ std) | $\text{RFG}_{\text{lin}}$ / $\text{RFG}_{\text{soft}}$ |
|---|---|---|---|---|---|---|
| **20% W/60% B/20% A** | Asian Female | 22 | 0.77 | 6.56 ± 5.62 | 6.37 ± 5.07 | |
| | Asian Male | 22 | 0.73 | 5.26 ± 6.74 | 5.61 ± 6.59 | |
| | Black Female | 27 | 0.67 | 7.49 ± 6.91 | 7.64 ± 6.63 | |
| | Black Male | 17 | 0.87 | 4.75 ± 4.59 | 4.80 ± 4.86 | |
| | White Female | 28 | 0.80 | 6.50 ± 4.53 | 5.89 ± 4.17 | |
| | White Male | 16 | 0.84 | 5.23 ± 5.09 | 5.38 ± 4.74 | 2.74 / 2.84 |
| **60% W/20% B/20% A** | Asian Female | 22 | 0.75 | 7.07 ± 5.70 | 6.35 ± 5.63 | |
| | Asian Male | 22 | 0.64 | 6.44 ± 7.51 | 6.58 ± 7.27 | |
| | Black Female | 27 | 0.55 | 9.27 ± 7.26 | 8.99 ± 6.80 | |
| | Black Male | 17 | 0.73 | 7.57 ± 6.69 | 7.57 ± 7.19 | |
| | White Female | 28 | 0.85 | 5.29 ± 4.79 | 5.23 ± 4.42 | |
| | White Male | 16 | 0.74 | 7.10 ± 6.00 | 6.68 ± 5.48 | 3.99 / 3.76 |
| **50% W/50% B** | Asian Female | 22 | 0.68 | 8.06 ± 6.74 | 6.96 ± 5.52 | |
| | Asian Male | 22 | 0.65 | 5.66 ± 7.63 | 6.47 ± 7.60 | |
| | Black Female | 27 | 0.63 | 8.11 ± 6.73 | 7.55 ± 6.86 | |
| | Black Male | 17 | 0.78 | 6.40 ± 6.38 | 5.71 ± 6.48 | |
| | White Female | 28 | 0.81 | 6.40 ± 4.78 | 5.64 ± 4.18 | |
| | White Male | 16 | 0.83 | 5.92 ± 4.07 | 5.81 ± 4.19 | 2.45 / 1.91 |

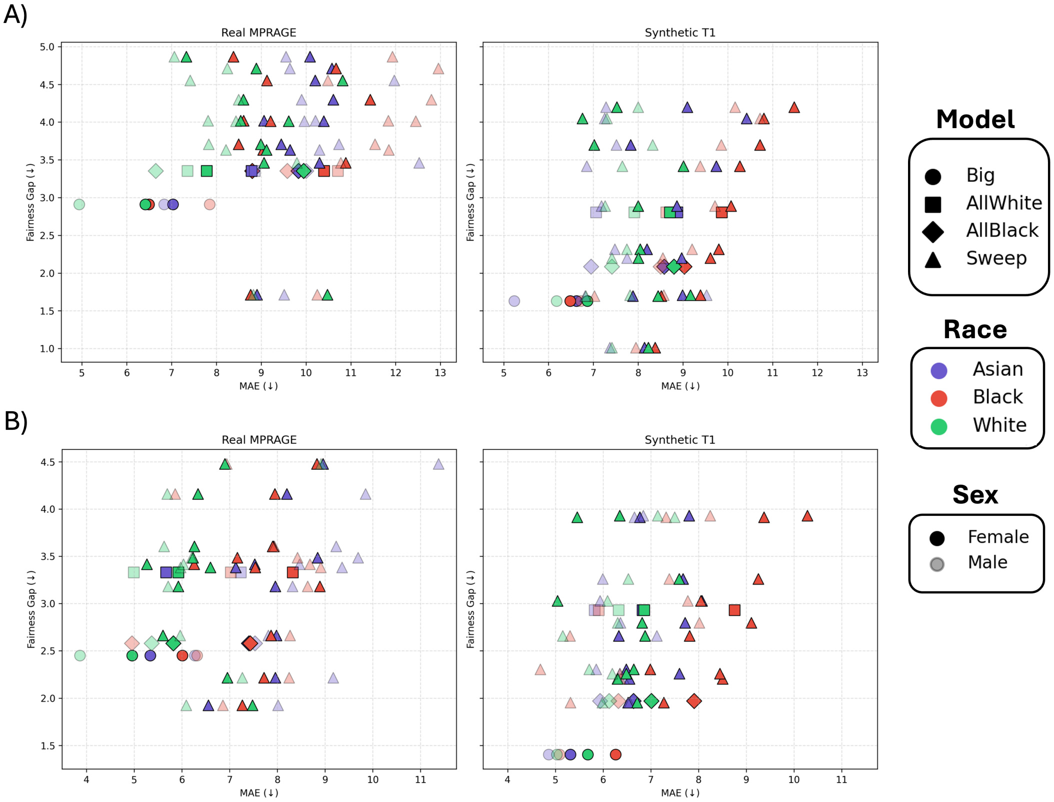


**Figure S4:** Relationship between subgroup mean absolute error and demographic fairness gap across all training configurations in the real T1-weighted and SuperSynth-derived synthetic T1 domains. Colors indicate racial group (Asian, Black, White), marker transparency indicates sex (Female, Male), and marker shape denotes model family (all-data, all-White, all-Black, and composition-sweep configurations). The fairness gap was computed across the six race-by-sex subgroups (Asian Female, Asian Male, Black Female, Black Male, White Female, White Male). (A) Beheshti-corrected results. (B) Final-corrected results after soft-Gaussian age-bin correction. Lower values on both axes indicate better performance and greater fairness.

### S7 External validation on ADNI3

#### Age-bias correction in ADNI3

The Beheshti correction was estimated using 5-fold cross-validation within the CN subset: coefficients were fitted on four folds and applied to the held-out fold, cycling until all CN subjects had out-of-fold corrected predictions. The resulting correction was then re-fitted on the full CN subset and applied to subjects with SMC and MCI. The soft-Gaussian age-bin correction was subsequently estimated from the Beheshti-corrected CN predictions and applied to all ADNI participants.

#### Results

**Table S9:** External validation on the ADNI3 dataset. Overall MAE (all clinical groups pooled) and CN-only $R^2$ and Pearson $r$ across the three correction stages. Bold values indicate the best-performing model within each real/synthetic training-domain comparison.

| | | Uncorrected | | | Beheshti | | | Fully corrected | | |
|---|---|---|---|---|---|---|---|---|---|---|
| **Model** | **Domain** | **MAE** | $R^2_{\text{CN}}$ | $r_{\text{CN}}$ | **MAE** | $R^2_{\text{CN}}$ | $r_{\text{CN}}$ | **MAE** | $R^2_{\text{CN}}$ | $r_{\text{CN}}$ |
| All-data | Real | $8.03 \pm 5.09$ | $-2.329$ | 0.682 | $4.61 \pm 3.59$ | 0.275 | 0.759 | $4.59 \pm 3.50$ | 0.269 | 0.781 |
| All-data | Synthetic | $\mathbf{7.27 \pm 4.40}$ | $-1.728$ | **0.737** | $\mathbf{4.18 \pm 3.40}$ | **0.393** | 0.788 | $4.17 \pm 3.37$ | 0.390 | 0.795 |
| 20% W/60% B/20% A | Real | $11.41 \pm 6.99$ | $-5.268$ | 0.530 | $5.94 \pm 4.51$ | $-0.307$ | 0.656 | $5.88 \pm 4.45$ | $-0.305$ | 0.681 |
| 20% W/60% B/20% A | Synthetic | $\mathbf{6.64 \pm 4.49}$ | $-1.457$ | **0.624** | $\mathbf{4.46 \pm 3.23}$ | **0.270** | 0.758 | $4.47 \pm 3.22$ | 0.270 | 0.681 |

*Note*: All $r$ values significant at $p < 0.0001$. $R^2_{\text{CN}}$ and $r_{\text{CN}}$ computed on CN subjects only ($n = 121$; predicted vs chronological age).

**Table S10:** ADNI3 clinical-group performance and disease-group comparisons at the fully corrected stage for the three calibrated combinations of model and imaging domain. CN rows report raw BAG (mean ± SEM, 95% CI); SMC and MCI rows additionally report the difference from CN (Welch's $t$-test: mean difference = group BAG − CN BAG, $t$, $p$, Cohen's $d$).

| | | | | | | | Difference from CN | | | |
|---|---|---|---|---|---|---|---|---|---|---|
| **Model** | **Domain** | **Group** | $N$ | **MAE** (± SD) | **BAG mean** (± SEM) | **BAG 95% CI** | **Diff.** (yr) | $t$ | $p$ | $d$ |
| All-data | Real | CN | 121 | $3.99 \pm 3.14$ | $+0.11 \pm 0.46$ | $[-0.80, +1.03]$ | — | — | — | — |
| | | SMC | 49 | $4.18 \pm 3.26$ | $+0.48 \pm 0.76$ | $[-1.04, +2.01]$ | 0.37 | 0.42 | 0.678 | 0.07 |
| | | MCI | 132 | $5.24 \pm 3.78$ | $+3.06 \pm 0.50$ | $[+2.08, +4.04]$ | 2.95 | 4.35 | $\mathbf{< 0.001}$ | 0.54 |
| All-data | Synthetic | CN | 121 | $3.43 \pm 3.10$ | $+0.02 \pm 0.42$ | $[-0.81, +0.86]$ | — | — | — | — |
| | | SMC | 49 | $3.68 \pm 3.22$ | $+0.99 \pm 0.69$ | $[-0.39, +2.37]$ | 0.96 | 1.20 | 0.235 | 0.21 |
| | | MCI | 132 | $4.90 \pm 3.46$ | $+3.14 \pm 0.45$ | $[+2.25, +4.02]$ | 3.11 | 5.07 | $\mathbf{< 0.001}$ | 0.64 |
| 20% W/60% B/20% A | Synthetic | CN | 121 | $3.83 \pm 3.24$ | $+0.01 \pm 0.46$ | $[-0.89, +0.92]$ | — | — | — | — |
| | | SMC | 49 | $4.18 \pm 2.86$ | $+1.58 \pm 0.69$ | $[+0.18, +2.97]$ | 1.56 | 1.89 | 0.062 | 0.31 |
| | | MCI | 132 | $5.00 \pm 3.29$ | $+2.35 \pm 0.48$ | $[+1.40, +3.30]$ | 2.34 | 3.53 | $\mathbf{< 0.001}$ | 0.44 |

### S8 Race-label injection implementation

#### Architecture

The race-injected variant of the SFCN concatenates the subject's one-hot race encoding to the globally pooled bottleneck feature vector immediately before the prediction head. The convolutional pathway is left entirely unchanged as race enters the network only after global average pooling, so any fairness benefit must arise from a learned race-specific linear correction at the output layer rather than from altered feature extraction.

#### Results

After training on the 60% W/20% B/20% A real-domain configuration, the model learned race-specific prediction offsets (estimated from mean predictions on the balanced test set) of approximately Asian: +3.46 years, Black: −1.63 years, White: −1.83 years, relative to the grand mean prediction. The Black–White offset difference was only 0.20 years, negligible given an observed Black–White MAE gap exceeding 2 years in the same configuration without injection. The model did not learn a targeted correction for Black prediction error; instead it learned a near-uniform shift nearly identical for Black and White subjects, alongside a spurious positive offset for Asian subjects likely reflecting an age-distribution imbalance in the training data. Integrated gradients attribution maps for the race-injected model were indistinguishable from those of the standard model (mean absolute attribution difference $6.95 \times 10^{-5}$; per-race differences 5.8–6.2%, not distinguishable from within-model noise), confirming that race supervision was absorbed entirely into the linear prediction head as a global intercept correction rather than reorganizing the convolutional feature space.

### S9 Representational analysis

#### Feature extraction

Representational analyses were performed on trained SFCN models with all network weights frozen. For each subject, representations were extracted from the output of each of the five convolutional blocks and from the final embedding layer immediately preceding the regression head. Feature maps were global-average-pooled across the spatial dimensions, yielding one feature vector per subject at each network depth; pooled dimensionality was 32, 64, 128, 256, and 256 for blocks 1–5 respectively, and 64 for the embedding layer. These pooled vectors formed the common input to both the $t$-SNE and the linear probing analyses.

Probes were trained on validation-set representations and evaluated on held-out test-set representations, both fixed across all experiments (132 participants each, 44 per racial group). Features were standardized using statistics estimated on the validation set only and applied unchanged to the test set.

$t$-SNE was applied to the best-fairness (20% W/60% B/20% A) and worst-fairness (60% W/ 20% B/20% A) configurations in both imaging domains; linear probing was additionally applied to the all-data and hybrid oversampling models.

#### Race probe

Race was decoded using multinomial logistic regression with an L2 penalty. The penalty strength $C$ was selected by 5-fold stratified cross-validation on the validation set over the grid $C \in \{0.01, 0.1, 1, 10\}$, scored by balanced accuracy, and the winning estimator was refitted on the full validation set before evaluation on the test set. Performance is reported as balanced accuracy against a chance level of 1/3, together with per-class accuracy and macro-averaged one-versus-rest AUC.

Significance was assessed by permutation testing: race labels were shuffled 1,000 times and the cross-validated balanced accuracy recomputed on each permutation, yielding an empirical null distribution. Permutation testing was performed on the validation set, so that the test set contributed only to the reported accuracies and never to model selection or inference. The resulting $p$-value therefore indicates whether race is decodable from a given representation, rather than whether the specific test-set accuracy differs from chance.

#### Age probe

Age was decoded using ridge regression, with the penalty $\alpha$ selected by 5-fold cross-validation on the validation set over 25 logarithmically spaced values from $10^{-3}$ to $10^{3}$. Performance is reported as the held-out coefficient of determination on the test set.

Regularization is necessary at this scale. The pooled feature dimension reaches 256 with only 132 validation subjects, making the design matrix rank-deficient at blocks 4 and 5. In these cases, the OLS coefficient vector is not uniquely identifiable; scikit-learn returns a minimum-norm least-squares solution. Even where the matrix is full rank, it can remain poorly conditioned. At block 3 the condition number of the scaled validation matrix exceeds $8 \times 10^3$ in the real domain and $9 \times 10^3$ in the synthetic domain, and held-out $R^2$ falls to $-17.5$ and $-3.2$ respectively. Across all 12 cells, unregularized least squares gives negative held-out $R^2$ in five (Table S11).

Cross-validation of $\alpha$ is likewise not cosmetic: the selected value ranged from 0.56 to $10^3$ across layers, spanning more than three orders of magnitude, so no single fixed penalty is appropriate at every depth. Using the `scikit-learn` default of $\alpha = 1$ instead cost on average 0.066 in held-out $R^2$, rising to 0.257 at block 5 in the real domain.

As for race, significance was assessed by permutation testing. The penalty was fixed at the cross-validated optimum $\alpha^\star$ and age labels were shuffled 1,000 times, with the cross-validated $R^2$ recomputed on each permutation to build an empirical null; re-selecting $\alpha$ within every permutation gives an equivalent $p$-value at roughly a hundred times the computational cost. Unlike balanced accuracy, $R^2$ has no lower bound, and the permutation null is accordingly negative.

Critically, the domain effect reported in the main text is unchanged under all three specifications: synthetic-domain representations yielded higher age $R^2$ than real-domain representations at all six depths under ordinary least squares, under ridge at fixed $\alpha = 1$, and under cross-validated ridge alike. The choice of estimator therefore affects the absolute decodability values but not the comparison the analysis is used to support.

**Table S11:** Age-probe estimator comparison across network depth and imaging domain. $p$ is the pooled feature dimension against $n = 132$ validation subjects; "rank" is the numerical rank of the scaled validation matrix, and ordinary least squares coefficient vector is not unique where rank $< p$. $R^2$ values are computed on the held-out test set. $\alpha_{\text{CV}}$ is the penalty selected by 5-fold cross-validation.

| Domain | Layer | $p$ | rank | cond($X$) | $R^2_{\text{OLS}}$ | $R^2_{\alpha=1}$ | $R^2_{\text{CV}}$ | $\alpha_{\text{CV}}$ |
|---|---|---|---|---|---|---|---|---|
| Real | block1 | 32 | 32 | 729 | −0.274 | −0.001 | −0.013 | 1000 |
| Real | block2 | 64 | 64 | 1167 | −0.103 | 0.211 | 0.214 | 0.56 |
| Real | block3 | 128 | 128 | 8474 | −17.481 | 0.235 | 0.320 | 5.62 |
| Real | block4 | 256 | 131 | 337 | −0.426 | 0.167 | 0.424 | 56.2 |
| Real | block5 | 256 | 131 | 198 | 0.042 | 0.344 | 0.587 | 56.2 |
| Real | embedding | 64 | 64 | 133 | 0.455 | 0.545 | 0.613 | 31.6 |
| Synth | block1 | 32 | 32 | 790 | 0.492 | 0.464 | 0.468 | 0.56 |
| Synth | block2 | 64 | 64 | 756 | 0.094 | 0.527 | 0.527 | 1.00 |
| Synth | block3 | 128 | 128 | 9670 | −3.181 | 0.524 | 0.544 | 1.78 |
| Synth | block4 | 256 | 131 | 292 | 0.388 | 0.539 | 0.575 | 17.8 |
| Synth | block5 | 256 | 131 | 155 | 0.598 | 0.671 | 0.733 | 56.2 |
| Synth | embedding | 64 | 63 | 209 | 0.644 | 0.736 | 0.761 | 178 |

Synthetic-domain $R^2$ exceeds real-domain $R^2$ at every depth under all three estimators.

#### $t$-SNE

$t$-SNE embeddings were computed independently at each network depth using a perplexity of 30 and 1,000 optimization iterations, and were colored by age, race, and sex to assess whether any of those variables produced visible gradients or cluster structure in the two-dimensional embedding.

Across configurations and domains, embeddings were organized primarily along an age gradient, with no clear global separation by race or sex (Figure S5). This is not in tension with the probing result: race was linearly decodable above chance from the same representations, indicating that demographic information is distributed across the representation.

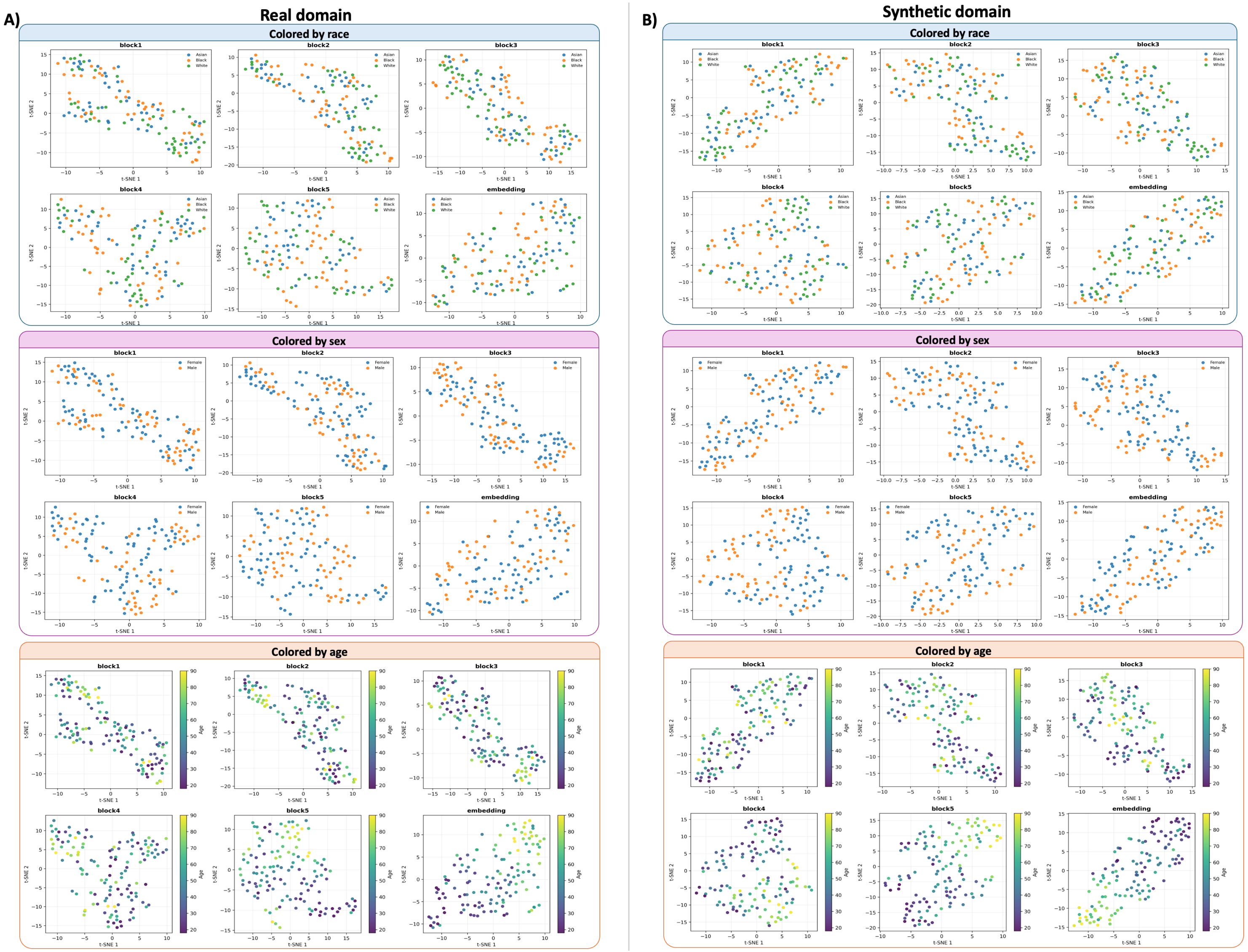


**Figure S5:** $t$-SNE embeddings of globally pooled representations at each network depth, colored by age, race, and sex. Perplexity 30, 1,000 optimization iterations.

### S10 Integrated gradients

#### Methods

Integrated gradients attribution maps (Sundararajan et al., 2017) were generated for each subject using a zero-image baseline and 50 integration steps. Attribution-difference maps were summarized using the SuperSynth segmentation of the MNI template: within each ROI, the signed mean difference quantified the direction of the attribution shift, while the mean absolute difference quantified its magnitude irrespective of direction. ROIs containing fewer than 500 voxels were excluded.

Two complementary ROI analyses were performed. The first was restricted to intracranial regions, including brain tissue, ventricles, and CSF spaces. The second retained all eligible ROIs, including vascular, ocular, muscular, skeletal, and other extracerebral tissues, in order to assess non-brain contributions to the attribution maps.

#### Results

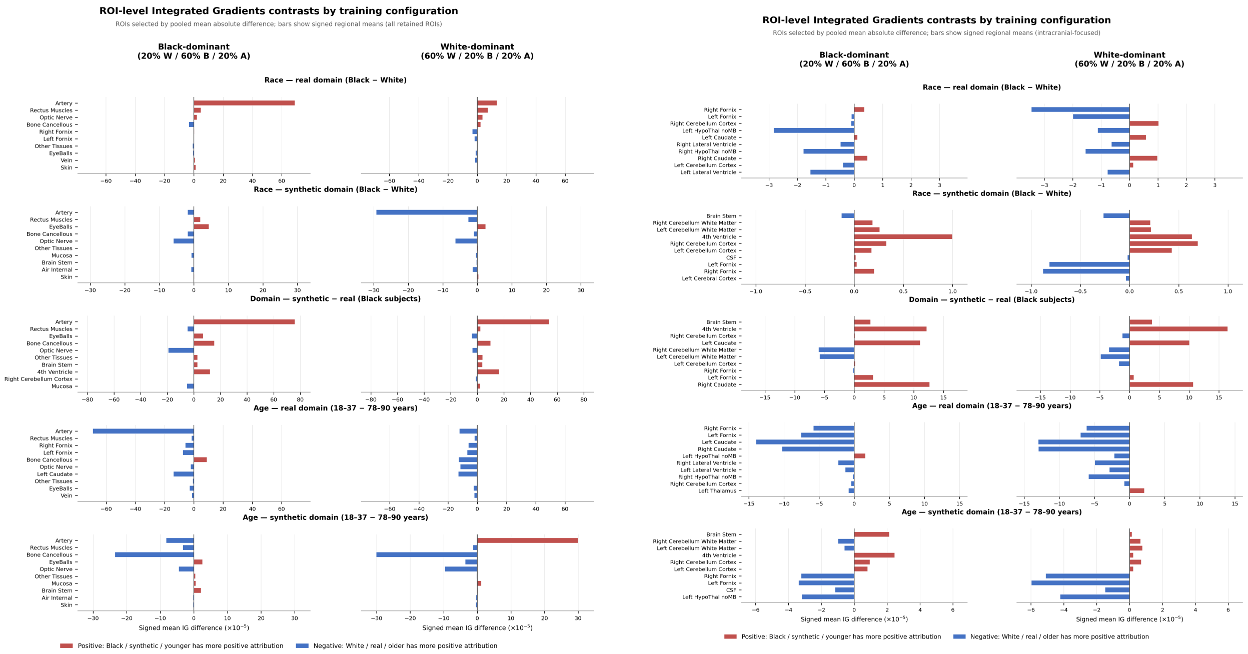


**Figure S6:** Left: all eligible ROIs, including extracerebral tissues. Right: intracranial ROIs, including brain tissue, ventricles, and CSF spaces.

Full per-ROI signed and absolute attribution differences for all three contrasts, in both configurations and both imaging domains, are shown in Figure S6. In the synthetic domain, age-related differences became more composition-dependent: the Black-dominant model showed older-positive differences in the hippocampus, amygdala, and hypothalamus, whereas the White-dominant model showed them predominantly in the hypothalamus, fornix, and third ventricle, with younger-positive attribution concentrated in the cerebellum and brainstem. The White-dominant model additionally showed a stronger real-positive thalamic difference, and the Black-dominant model a stronger real-positive hypothalamic difference.

#### References

Ioffe, S. & Szegedy, C. Batch normalization: Accelerating deep network training by reducing internal covariate shift. In: *Proceedings of the 32nd International Conference on Machine Learning (ICML)*. Vol. 37*, 2015*, pp. 448–456.

Sundararajan, M., Taly, A. & Yan, Q. (2017) *Axiomatic attribution for deep networks.* https://arxiv.org/abs/1703.01365

Wu, Y. & He, K. Group normalization. In: *Computer Vision – ECCV 2018, 2018.* : Springer.
